# CUX1 and IκBζ mediate the synergistic inflammatory response to TNF and IL-17A in stromal fibroblasts

**DOI:** 10.1101/571315

**Authors:** Kamil Slowikowski, Hung N. Nguyen, Erika H. Noss, Daimon P. Simmons, Fumitaka Mizoguchi, Gerald F.M. Watts, Michael F. Gurish, Michael B. Brenner, Soumya Raychaudhuri

## Abstract

The role of stromal fibroblasts in chronic inflammation is unfolding. In rheumatoid arthritis (RA), leukocyte-derived cytokines tumor necrosis factor (TNF) and IL-17A work together, activating fibroblasts to become a dominant source of the hallmark cytokine IL-6. However, IL-17A alone has minimal effect on fibroblasts. To identify key mediators of the synergistic response to TNF and IL-17A in human synovial fibroblasts, we performed time series, dose response, and gene silencing transcriptomics experiments. Here we show that in combination with TNF, IL-17A selectively induces a specific set of genes mediated by factors including CUX1 and IκBζ. In the promoters of *CXCL1*, *CXCL2*, and *CXCL3*, we found a putative CUX1-NF-κB binding motif not found elsewhere in the genome. CUX1 and NF-κB p65 mediate transcription of these genes independent of LIFR, STAT3, STAT4, and ELF3. Transcription of *NFKBIZ*, encoding the atypical IκB factor IκBζ, is IL-17A dose-dependent, and IκBζ only mediates the transcriptional response to TNF and IL-17A, but not to TNF alone. In fibroblasts, IL-17A response depends on CUX1 and IκBζ to engage the NF-κB complex to produce chemoattractants for neutrophil and monocyte recruitment.

## Introduction

In rheumatoid arthritis (RA), stromal fibroblasts are key components of the hyperplastic and invasive synovial tissue that mediates joint destruction and bone resorption (Bottini and Firestein, 2013). Interleukin 17 (IL-17A), a cytokine produced by CD4^+^ Th17 cells (Chabaud et al., 1999) and other cells (Monin and Gaffen, 2018), is crucial to perpetuate the vicious inflammatory cycle localized to the joint tissue that contributes to induction, progression, and chronicity of RA (Benedetti and Miossec, 2014). Briefly, leukocytes activate stromal cells to produce inflammatory cytokines and chemokines such as IL-6, IL-8, CCL2 (Zrioual et al., 2009), and GM-CSF (Hirota et al., 2018). In turn, these cytokines attract more leukocytes to infiltrate the synovial tissue and activate them to produce more IL-17A. The infiltrating leukocytes produce tumor necrosis factor alpha (TNF), and IL-17A synergizes with TNF to further amplify the pathways underlying chronic inflammatory disease pathogenesis (Monin and Gaffen, 2018).

The combination of TNF and IL-17A synergistically activates synovial fibroblasts to produce IL-6 (Zrioual et al., 2009), and these cells are a dominant source of IL-6 in the joint tissue (Nguyen et al., 2017; Zhang et al., 2018). While other cytokines are involved in RA, clinical efficacy of targeting TNF and IL-6 has clearly established their pivotal roles in pathogenesis (Smolen and Aletaha, 2015). Clinical relevance of the synergistic activity of TNF and IL-17A has also been shown: measurements of TNF and IL-17A mRNA in RA synovial tissue predict early joint damage progression by magnetic resonance imaging (MRI) (Kirkham et al., 2006). Joint damage progresses as TNF and IL-17A induce synovial fibroblasts to produce matrix metalloproteinases (MMPs) for digesting extracellular matrix and invading the cartilage (Moran et al., 2009). In addition, IL-17A plays a role in differentiation of osteoclast progenitors into bone-resorbing osteoclasts via increased expression of RANKL in osteoblasts (Kotake et al., 1999).

However, most studies of IL-17A transcriptional response have used qPCR to measure only the most critical genes with known roles in the context of inflammation, osteoclastogenesis, angiogenesis, or neutrophil recruitment (Chabaud et al., 2001; Ermann et al., 2014; Fossiez et al., 1996; Koshy et al., 2002; Moran et al., 2009; Ruddy et al., 2004; Shen et al., 2005; Zrioual et al., 2009). One study used microarrays to measure genome-wide gene expression response in a single sample 12 hours after stimulation with IL-17A and IL-17F alone or in combination with TNF (Zrioual et al., 2009). An expanded view with robust measurements of transcriptional dynamics over time would help to reveal mediators in regulatory pathways downstream of TNF and IL-17A. While these regulatory pathways are well studied in leukocytes (Lubberts, 2015), there remains much to be learned about the transcriptional regulation of inflammatory functions in mesenchymal cells such as synovial fibroblasts. For example, we recently described an autocrine loop driving IL-6 production that is only used by fibroblasts, and not by leukocytes, involving LIF, LIFR, and STAT4 (Nguyen et al., 2017). Many different cell types have been shown to use the NF-κB signaling pathway for inflammatory response, but additional components of this pathway are being discovered that modulate cell type and condition-specific responses to inflammatory signals (Zhang et al., 2017).

Surprisingly, IL-17A in the absence of TNF has a minimal transcriptional effect in fibroblasts, and defining its role with TNF requires a highly sensitive study. To identify key mediators of the response to TNF and IL-17A in human synovial fibroblasts, we use transcriptomics data from a high-resolution time series with multiple IL-17A doses. By measuring the *in vitro* transcriptomic effects caused by stimulation with TNF or costimulation with TNF and IL-17A at three doses of IL-17A (0, 1, 10 ng/mL), we identify genes with transcriptional changes proportional to IL-17A dose. We also find that transcription of specific sets of chemokines is controlled by different mediators. A molecular complex of NF-κB p65 and CUX1 mediates transcription of CXC chemokines (*CXCL1*, *CXCL2*, *CXCL3*, *CXCL6*). In contrast, LIFR mediates transcription of CC chemokines (*CCL7*, *CCL8*, *CCL20*). We distinguish these transcriptional responses by silencing each mediator with short interfering RNAs (siRNAs) and then assaying the downstream effects. Furthermore, we find that transcription of atypical IκB factor IκBζ (NFKBIZ) is responsive to IL-17A dose, and IκBζ mediates IL-6, IL-8, and MMP-3 production only after costimulation with TNF and IL-17A, but not after TNF alone. Finally, we demonstrate that fibroblasts require these mediators to recruit neutrophils and monocytes in an *in vitro* assay.

## Results

### TNF and IL-17A synergistically activate IL-6 and MMP-3 production

To demonstrate the synergy between TNF and IL-17A, we stimulated synovial fibroblasts with 16 combinations of dosages of TNF (0, 0.1, 1, 10 ng/mL) and IL-17A (0, 1, 10, 100 ng/mL). We cultured primary fibroblasts from two RA donors and measured IL-6 protein levels in the supernatant after stimulation by enzyme-linked immunosorbent assay (ELISA). Stimulation with one cytokine (TNF or IL-17A) is significantly less effective at inducing IL-6 than costimulation with TNF and IL-17A. We saw 1.2 ng/mL IL-6 after stimulation with IL-17A alone (100 ng/mL), and 2.6 ng/mL IL-6 after TNF alone (10 ng/mL) (**Figure 1A**). In contrast, we saw 49.3 ng/mL of IL-6 in the supernatant after costimulation with TNF (10 ng/mL) and IL-17A (100 ng/mL) – the addition of IL-17A caused a 20-fold increase in IL-6 levels relative to stimulation with TNF alone (**Figure 1A**). Matrix Metalloproteinase 3 (MMP-3) protein levels showed a similar synergistic response to TNF and IL-17A (**Figure 1A**). These results confirm previous reports that TNF and IL-17A synergistically activate IL-6, MMP-3, and other inflammatory cytokines (Chabaud et al., 2001; Fossiez et al., 1996; Koshy et al., 2002; Moran et al., 2009; Ruddy et al., 2004; Shen et al., 2005).

**Figure 1.**
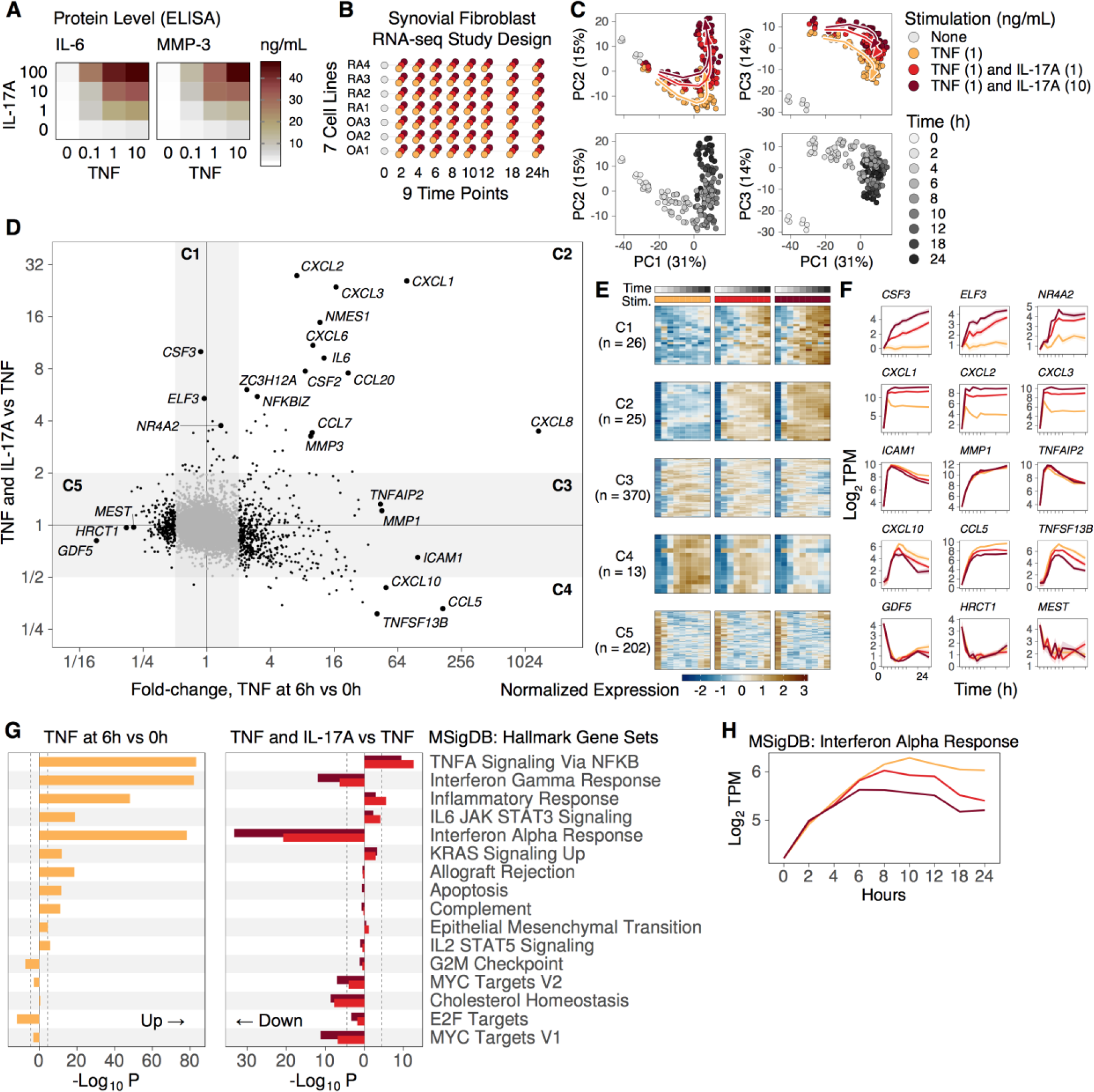
Defining the transcriptional response to TNF and IL-17A in fibroblasts with a high resolution time series and dose response study. **(A)** Protein levels of IL-6 and MMP-3 measured by ELISA after stimulating with 16 different combinations of doses: TNF (0, 0.1, 1, 10 ng/mL) and IL-17A (0, 1, 10, 100 ng/mL). Costimulation with TNF and IL-17A results in synergistic IL-6 and MMP-3 protein production. **(B)** Our study design includes 7 cell lines from patient tissues. We stimulated them with TNF (1 ng/mL) and three doses of IL-17A (0, 1, and 10 ng/mL). We measured gene expression at baseline and 8 time points after stimulation. **(C)** Principal Component Analysis (PCA) with 636 genes responsive to TNF alone or TNF and IL-17A. Samples fall along the principal curves. 60% of the total variation is captured by the first 3 principal components. **(D)** Differential expression analysis shows that genes fall into 5 major categories, depending on the response to TNF alone (x-axis) or to TNF and IL-17A (y-axis). **(E)** Heatmaps with normalized expression for 636 genes, where rows are genes and columns are time points. From left to right, each heatmap shows response to TNF (1 ng/mL) and each of the three doses of IL-17A (0, 1, or 10 ng/mL). **(F)** Mean gene expression (lines) and SEM (ribbons) over time for genes with characteristic profiles: *CSF3, ELF3, NR4A2, CXCL1, CXCL2, CXCL3, ICAM1, MMP1, TNFAIP2, CXCL10, CCL5, TNFSF13B, GDF5, HRCT1, MEST.* **(G)** Gene set enrichment with MSigDB Hallmark gene sets. The x-axis shows signed p-values indicating the strength of enrichment with induced or repressed genes for TNF at 6h versus 0h (left) and TNF and IL-17A (1 or 10 ng/mL) versus TNF (right). **(H)** We selected the 10 genes in the Interferon Alpha Response gene set with the most significant p-values for TNF and IL-17A (10 ng/mL) versus TNF, and show mean +/-SEM of the means of the 10 genes.

### IL-17A requires TNF to induce a gene expression response in fibroblasts

Since IL-17A resulted in synergistic protein production of IL-6 and MMP-3, we decided to assay the direct genome-wide transcriptional effects of IL-17A. To do this, we conducted two initial genome-wide transcriptional experiments after stimulation with IL-17A or TNF, alone or in combination.

First, we profiled one synovial fibroblast cell line at 9 time points over 72 hours in 4 different conditions: media only, TNF (1 ng/mL), IL-17A (1 ng/mL), and TNF with IL-17A (1 ng/mL and 1 ng/mL). We used microarrays to assay the transcriptional response (**Microarray Data 1**, **Figure S1**, **Table S1**). After stringent quality control, we fit a linear model to each gene to find which genes show differential expression between conditions (**Methods**). In response to IL-17A alone, 3 genes (*NFKBIZ*, *TNFAIP6*, and *CXCL1*) were differentially expressed relative to media (FC > 1.5 and FDR < 0.05, **Figure S1D**). On the other hand, 1,208 genes responded to TNF alone (**Figure S1E**). In response to TNF and IL-17A, 15 genes were differentially expressed relative to TNF alone (**Figure S1F**). We also applied principal components analysis (PCA) (Raychaudhuri et al., 2000) to **Microarray Data 1**. Along the first two components, samples stimulated with IL-17A alone are most similar to unstimulated samples, and there is little difference between samples stimulated with TNF alone or TNF and IL-17A (**Figure S1B,C**). We conclude that that IL-17A is dependent on the presence of TNF to induce significant effects on gene expression, consistent with our ELISA results for protein levels of IL-6 and MMP-3 (**Figure 1A**).

Given the subtle transcriptional difference between costimulation with TNF and IL-17A versus TNF alone, we ran a second experiment where we assayed gene expression response in four synovial fibroblast cell lines (2 RA and 2 OA) 24 hours after stimulation (**Microarray Data 2**, **Figure S2**, **Table S2**). This experiment replicated the differential expression signal between TNF and IL-17A versus TNF from Microarray Data 1 (r = 0.67, P < 10^-15^). This gave us confidence that a well-powered transcriptional study could identify the downstream mediators that respond to costimulation with TNF and IL-17A.

### Defining the transcriptional response to TNF and IL-17A in fibroblasts with a high resolution time series and dose response study

We used RNA-seq to assay transcriptomes of 7 synovial fibroblast cell lines (4 RA, 3 OA) at 9 time points over 24 hours after stimulation with TNF (1 ng/mL) and IL-17A (0, 1, and 10 ng/mL), resulting in a total of 175 RNA-seq profiles (**Figure 1B**, **Methods**). By using RA and OA cell lines, we could look for responses consistent across tissue states. By using multiple IL-17A dosages, we could identify genes with expression proportional to IL-17A dose. By modeling the dynamics of each gene over time, we increased our statistical power to identify differentially expressed genes (**Methods**).

Broadly, we found that genes could be subdivided into categories based on their response to TNF and IL-17A. We used linear models to estimate the effect sizes for each gene (see **Methods, Table S3**). Next, we identified the genes with greatest response to (1) TNF at 6h vs 0h and (2) costimulation with TNF and IL-17A vs TNF alone (**Figure 1C**). We limited our analysis to the 636 statistically significant genes (absolute FC > 2 and FDR < 0.05) and subdivided them into 5 categories (**Figure 1D**). We also applied principal components analysis (PCA) (Raychaudhuri et al., 2000) to these 636 genes (**Figure 1E**, **Figure S3**), and found that the first component (explaining 31% of the total variance) identifies genes with strong activation or deactivation over time in response to TNF stimulation. The second (15% of variance) and third (14% of variance) identify genes with expression altered by IL-17A dose.

### Transcriptional response reveals genes selectively induced by TNF and IL-17A costimulation or TNF alone

We defined Category C1 as the set of genes induced by costimulation with TNF and IL-17A, but unchanged by TNF alone (**Figure 1D,F**). C1 includes 26 genes with dose-dependency on IL-17A such as *CSF3*, encoding granulocyte colony stimulating factor G-CSF, a crucial factor for neutrophil survival and development (Panopoulos and Watowich, 2008; Parsonage et al., 2008) (**Figure S4**, **Table S3**). Interleukin 11 (*IL11*), a cytokine that is elevated in RA and promotes fibroblast invasion (Elshabrawy et al., 2018), was also transcribed in an IL-17A dose-dependent manner. The transcription factor (TF) with greatest expression response to IL-17A was the epithelium-specific ETS factor encoded by *ELF3*, so we hypothesized that ELF3 might mediate the IL-17A dose-dependent expression of other genes. For this reason, we included it in an siRNA gene silencing experiment (see below). Another TF in Category C1 is the orphan nuclear receptor encoded by *NR4A2*, which has been shown to induce synoviocyte proliferation and invasion (Mix et al., 2012), and also to directly control expression of *CXCL8* in inflammatory arthritis (Aherne et al., 2009). Category C1 also includes “POU domain, class 2, transcription factor 2” encoded by *POU2F2*: a TF with a binding site motif in the promoters of *CSF3* and immunoglobulin genes (**Figure S4**).

Category C2 has 25 genes induced in response to TNF and further induced by addition of IL-17A (**Figure 1D,F**), including members of the CXC subfamily of chemokines *CXCL1*, *CXCL2*, *CXCL3*, *CXCL6*, and *CXCL8*. Notably, *CXCL8* mRNA was 1000-fold more abundant after TNF at 6h relative to 0h, and the other 4 chemokines were 17-fold more abundant on average (**Figure S4**, **Table S3**). While TNF alone caused increased CXC chemokine transcription, the addition of IL-17A further increased transcription of these five chemokines in a dose-dependent manner (an additional 7-fold and 15-fold for 1 and 10 ng/mL IL-17A, respectively) (**Figure S4**). We note that IL-8 (CXCL8), CXCL1, and CXCL2 are potent neutrophil chemoattractants, complementing the function of G-CSF (encoded by *CSF3*).

Category C3 genes increase in response to TNF, but they are unaffected by the addition of IL-17A (**Figure 1D,F**). This is the largest category with 370 genes, including TNF alpha induced genes *TNFAIP6* and *TNFAIP2*, as well as matrix metalloproteinase 1 (*MMP1*) and bradykinin receptor (*BDKRB1*) (**Figure S4**, **Table S3**). We tested MSigDB Hallmark gene sets (Liberzon et al., 2015) for enrichment with TNF responsive genes and found significant enrichments in the gene sets "TNFA signaling via NFKB", "Interferon Alpha Response", and "Interferon Gamma Response" (**Figure 1G**).

Category C4 genes are transcribed in response to TNF, but transcription is dampened by the addition of IL-17A. C4 includes 13 genes related to interferon response such as *IRF1, IRF9, IFI30,* and *CXCL11* (**Figure S5**). Gene set enrichment analysis shows that genes in the MSigDB hallmark gene set "Interferon Alpha Response" were induced by TNF, but repressed by TNF and IL-17A (**Figure 1G**). Indeed, an increase from 1 ng/mL IL-17A to 10 ng/mL Il-17A causes more repression of these genes (**Figure 1H**). This suggests that TNF activates – and IL-17A attenuates – the genes responsive to interferon alpha.

Category C5 has 202 genes repressed by TNF but largely unaffected by IL-17A. Enrichment with MSigDB hallmark pathways reveals that many of these genes are related to cell cycle such as *STMN1, CDKN2C, CDC20, CDKN3, HMGB2, CDK1, CCNB2, CDCA8, TOP2A* (**Figure S5**). While TNF stimulation caused reduced transcription of genes involved in cell division, it caused increased transcription of genes involved in apoptosis such as *SOD2* and *BIRC3*. The genes with greatest reduction after TNF stimulation include growth differentiation factor 5 (*GDF5*), histidine rich carboxyl terminus 1 (*HRCT1*), and mesoderm specific transcript (*MEST*) (**Figure 1D**). For example, *GDF5* – a critical factor for normal synovial joint function (Parrish et al., 2017) and synovial lining hyperplasia (Roelofs et al., 2017) – decreased 11-fold (fold change 0.09, 95% CI 0.058-0.14, P = 3e-20) after 6h of TNF.

We conclude that TNF alone is sufficient to activate transcription of the CXC subfamily of neutrophil recruitment chemokines, and that IL-17A synergizes with TNF to increase transcription of these chemokines. Costimulation with both TNF and IL-17A is required for transcription of *CSF3*, while TNF alone is insufficient. Thus, costimulation of synovial fibroblasts with TNF and IL-17A activates a transcriptional program for recruiting neutrophils with CXC chemokines and sustaining neutrophil survival with G-CSF (*CSF3*).

### Transcription factors underlying the synergy between TNF and IL-17A

We used two strategies to find putative transcription factors that mediate the response to costimulation with TNF and IL-17A. First, we identified TFs with transcription proportional to IL-17A dose (Categories C1 and C2), including *ELF3*, *POU2F2, ZC3H12A*. Since *ELF3* expression increased (FC 5.4, 95% CI 4.4-6.6, P = 2e-37) after stimulation with IL-17A (10 ng/mL), we decided to study the global transcriptome-wide effects of silencing *ELF3* with siRNA.

Next, we searched for TF binding motifs in the promoters of differentially expressed genes. We used HOMER (Heinz et al., 2010) to perform *de novo* motif enrichment analysis with 26 genes within Categories C1 and C2 that had greater expression for 10 versus 1 ng/mL IL-17A (log2 FC > 1.5 (FC > 2.83) and FDR < 0.05) (**Methods**). As expected, we found motifs that closely resemble NF-κB, REL, and JUND binding sites (**Table 1**). We also found a motif that might represent the binding site for cut like homeobox 1 (CUX1), TCCGGATCGATC, that is present in the promoters of 4 of the 26 genes (P = 1e-10): *CXCL1, CXCL2, CXCL3, PIM2*. When we examined public CUX1 ChIP-seq data from ENCODE for three cell lines (GM12878, K562, MCF-7), we found evidence that CUX1 might bind these motifs. Consistent with our motif analysis and differential expression analysis, there are CUX1 ChIP-seq peaks in the promoters of *CXCL2*, *CXCL3*, *C15orf48 (NMES1)*, and *IL6*, as well as less than 5kb upstream of the promoter of *CXCL6* (**Figure S6A**). These results motivated us to assay the transcriptome-wide effects of silencing *CUX1*.

**Table 1.** HOMER *de novo* motifs in gene promoters.

| Best Match | P | FG (%) | BG (%) | Consensus | Promoters |
| --- | --- | --- | --- | --- | --- |
| NFkB-p65-Rel(RHD)/ThioMac-LPS-Expression(GSE23622) | 10<sup>-12</sup> | 16 | 0.0043 | CAGGGAATTTCC | CXCL1, CXCL2, CXCL3, CXCL6 |
| CUX1/MA0754.1/Jaspar | 10<sup>-10</sup> | 16 | 0.016 | TCCGGATCGATC | CXCL1, CXCL2, CXCL3, PIM2 |
| Nur77(NR)/K562-NR4A1-ChIP-Seq(GSE31363) | 10<sup>-10</sup> | 16 | 0.025 | ACCTTYMCAWKA | CCL20, CSF2, MMP3, TNFSF13B |
| REL/MA0101.1/Jaspar | 10<sup>-9</sup> | 20 | 0.11 | TCCGGGSTTTCC | CXCL1, CXCL2, CXCL3, NFKBIZ, PTGS2 |
| GLI2/MA0734.1/Jaspar | 10<sup>-9</sup> | 16 | 0.027 | GACCGCGCTGGC | C15orf48, CD83, G0S2, IL4I1 |
| Hic1/MA0739.1/Jaspar | 10<sup>-8</sup> | 24 | 0.51 | GCGATGGCCC | C15orf48, CCL20, CXCL1, CXCL2, CXCL3, G0S2, NFKBIZ |
| EWSR1-FLI1/MA0149.1/Jaspar | 10<sup>-7</sup> | 12 | 0.019 | GAAGGAAGGCGA | CXCL1, CXCL2, CXCL3 |
| JUND/MA0491.1/Jaspar | 10<sup>-6</sup> | 20 | 0.56 | GACTCATCCT | CCL7, CSF2, IL11, MMP3, NFKBIZ |

HOMER *de novo* motif analysis of the promoters of 25 genes with statistically significant difference ( $\log_2$ FC > 1.5 and FDR < 0.05) for 10 versus 1 ng/mL of IL-17A. For each discovered motif, we show the name of a known motif that is similar to the newly discovered motif, the hypergeometric p-value from HOMER, the percent of matches in the foreground (FG) gene set of 25 genes, the percent of matches in the background (BG) gene set of 39,486 genes, and the list of genes in the foreground that have a match for the motif.

In addition to silencing *ELF3* and *CUX1*, we also decided to silence the genes encoding leukemia inhibitory factor receptor (*LIFR*) and signal transducer and activator of transcription 4 (*STAT4*). Previously, we found that LIFR and STAT4 are key components of an autocrine loop controlling IL-6 production that is specific to fibroblasts and not leukocytes (Nguyen et al., 2017). We also decided to silence *STAT3* because it encodes a known mediator of inflammatory pathways (Hillmer et al., 2016), and we wanted to see how its function might be distinct from STAT4 in the fibroblast response to inflammatory chemokines.

### Transcriptomics time series after silencing putative mediators

We selected five genes encoding putative mediators of IL-17A transcriptional effects (*ELF3, CUX1, LIFR, STAT3, STAT4*), silenced them with short interfering RNAs (siRNAs), and assayed global gene expression with RNA-seq. We used qPCR to confirm that each of our selected siRNAs silences its intended target gene (**Figure S7A**). After incubating 4 RA synovial fibroblast cell lines with a siRNA for 2 days, we stimulated them with TNF (0.1 ng/mL) alone or TNF and IL-17A (1 ng/mL) and assayed the transcriptomes at four time points (0, 1, 6, and 16 hours) (**Figure 2A**). This experimental design let us confirm that these putative mediators are acting downstream of IL-17A, and allows us to directly compare their transcriptional effects.

**Figure 2.**
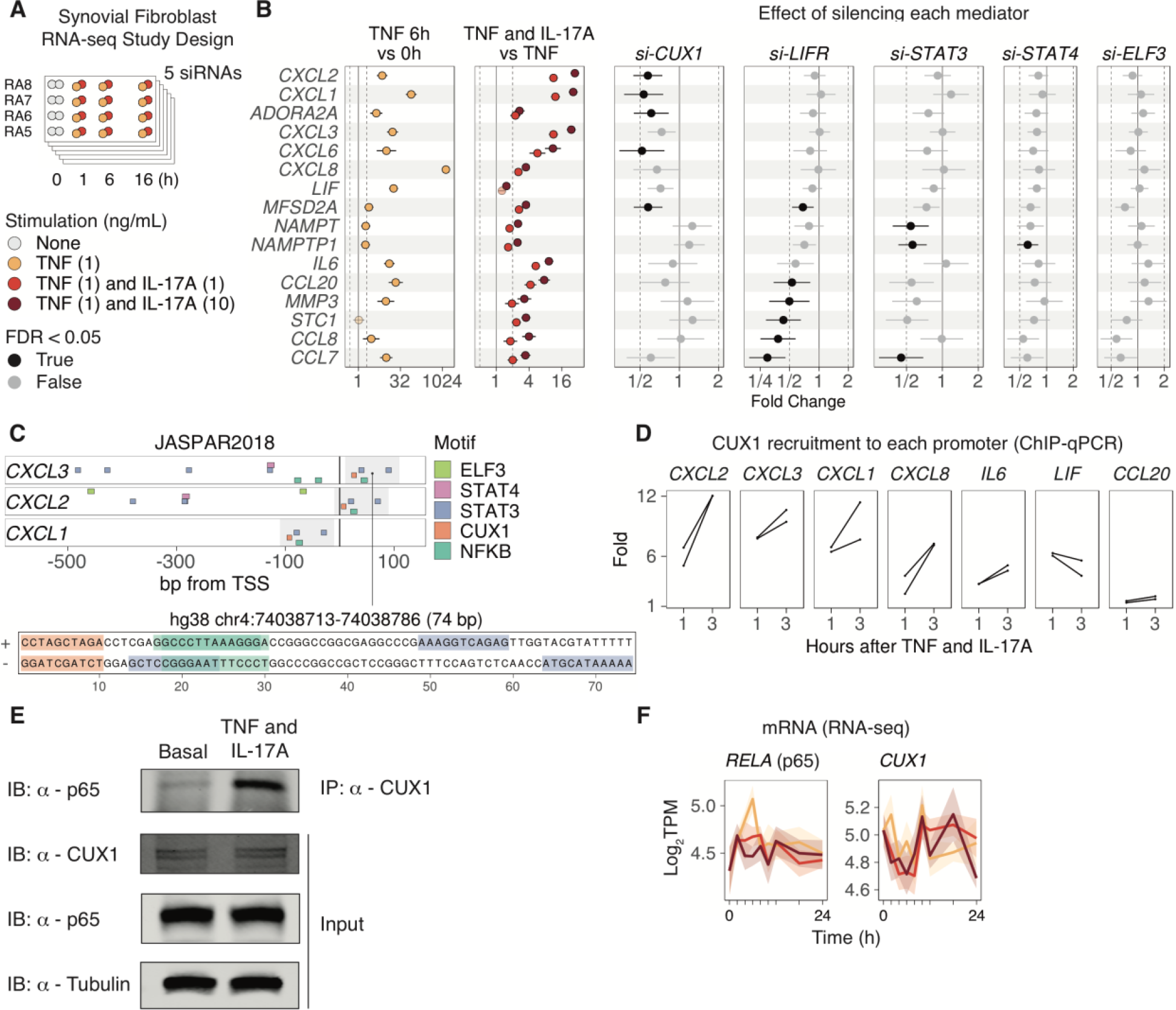
CUX1 mediates transcription of *CXCL1*, *CXCL2*, *CXCL3*, independent of LIFR, STAT3, STAT4, and ELF3. **(A)** Our experimental design included 4 synovial fibroblast cell lines from patient tissues, 4 time points (0, 1, 6 and 16 hours), 2 stimulations (TNF (0.1 ng/mL); TNF (0.1 ng/mL) and IL-17A (1 ng/mL)), and 5 siRNAs (*CUX1, LIFR, STAT3, STAT4, ELF3*). **(B)** Fold change estimates and 95% confidence intervals for 16 select genes that were measured by ChIP-qPCR, or that were responsive to IL-17A and had at least one significant siRNA effect. Left 2 panels show effect estimates from **RNA-seq Data 1**, and right 5 panels show effect estimates from **RNA-seq Data 2.** Dark color indicates FDR < 0.05. **(C)** The promoters of *CXCL1*, *CXCL2*, and *CXCL3* have a unique 74 bp sequence that contains motifs for CUX1, NF-κB, and STAT3. **(D)** ChIP-qPCR of the CUX1 protein to 7 gene promoters at 3 time points (0, 1, and 3 hours) after costimulation with TNF and IL-17A. The y-axis shows fold recruitment relative to the first time point at 0 hours, shown for 2 technical replicates. **(E)** After 4 hours of stimulation with TNF (1 ng/mL) and IL-17A (1 ng/mL), NF-κB p65 is detected in a pull down of CUX1. **(F)** The mRNA levels of *RELA* (p65) and *CUX1* are insensitive to IL-17A, and are relatively stable over time.

When we examined genes in Categories C1 and C2, we found two distinct groups of genes with different responses to gene silencing (**Figure 2B**). One group of genes was more repressed by silencing *CUX1*, and the other group was more repressed by silencing *LIFR*. Consistent with our motif analysis, silencing *CUX1* repressed Category C2 genes *CXCL1*, *CXCL2*, *CXCL3*, *CXCL6*, *CXCL8* (mean FC 0.60, 95% CI: 0.43-0.84). In contrast, these five genes were not strongly affected by silencing *ELF3*, *LIFR*, *STAT3*, or *STAT4* (mean FC 0.92, 95% CI: 0.66-1.3) (**Figure 2B**). We validated this result by using qPCR to measure selected genes in the same cDNA libraries (**Figure S7B**). These data suggest at least two independent regulatory pathways for genes induced by costimulation with TNF and IL-17A.

We downloaded the JASPAR2018 database of genome-wide transcription factor binding motif matches (Khan et al., 2018) to look more closely at gene promoters, and we found a unique CUX1-NF-κB nucleotide sequence motif in the promoters of *CXCL1, CXCL2, and CXCL3* (**Figure 2C**). Next, we ran a genome-wide BLAST search for the 74 bp sequence in the promoter of *CXCL3* (hg38 chr4:74038713-74038786) containing the CUX1 and NF-κB motifs. This sequence is present only in four genomic locations: the promoters of *CXCL1, CXCL2, CXCL3,* and pseudogene *CXCL1P1* (**Figure S6B**). Next, we used ChIP-qPCR to test if CUX1 binds selected promoters after stimulation with TNF and IL-17A. Indeed, CUX1 recruitment to the promoters of *CXCL1, CXCL2, CXCL3* increases more than 10-fold, on average, after 3 hours of stimulation relative to baseline (**Figure 2D**). Since CUX1 binds the promoters of genes with the CUX1-NF-κB motif, we hypothesized that CUX1 might bind NF-κB. Using an antibody against CUX1, we detected an increased amount of NF-κB p65 in the pull-down fraction after stimulating fibroblasts with TNF and IL-17A (**Figure 2E**). Since protein and mRNA levels of RELA and CUX1 remain relatively constant over time (**Figure 2E,F**), our data suggests that CUX1 and NF-κB form a molecular complex after costimulation with TNF and IL-17A. This complex likely binds the promoters of CXC chemokines and regulates transcription of these genes in synovial fibroblasts.

In addition to the group of genes repressed by silencing *CUX1*, a second group of genes was repressed after silencing *LIFR*. This includes the genes encoding C-C type chemokines encoded by *CCL8*, *CCL20*, as well as *MMP3* (Matrix metalloproteinase-3) and *STC1* (Stanniocalcin-1) (mean FC 0.34, 95% CI: 0.22-0.51, **Figure 2B**). These four genes were unaffected after silencing *CUX1* (mean FC 1.0, 95% CI: 0.67-1.6), suggesting they are not mediated by CUX1. CCL8 (MCP-2) is a chemoattractant and activator of monocytes, whereas CCL20 is a chemoattractant for lymphocytes and is known to recruit CCR6^+^ Th17 cells to sites of inflammation (Hirota et al., 2007). Our data suggests that two independent pathways activate in response to costimulation with TNF and IL-17A, and chemoattraction of leukocytes is a primary function of this response in synovial fibroblasts.

### IκBζ binds NF-κB p65 and p100/52, and mediates the response to TNF and IL-17A

Since NF-κB is a known mediator of inflammatory pathways, and *NFKBIZ* transcriptional response is driven by IL-17A dose (**Figure 3A**), we tested the effects of silencing *NFKBIZ*, the gene encoding IκBζ. We measured protein levels of IL-6, IL-8 (CXCL8), and MMP-3 after stimulating synovial fibroblasts with TNF alone or with the combination of TNF and IL-17A. Silencing *NFKBIZ* reduced protein levels of IL-6, IL-8, and MMP-3 after TNF and IL-17A, but not after TNF alone (**Figure 3B**). Next, we tested if IκBζ binds NF-κB to modulate the transcription of its downstream targets in fibroblasts. Using an antibody against NF-κB p65, we detected increased levels of IκBζ in the pull-down fraction after stimulating fibroblasts with TNF and IL-17A relative to unstimulated fibroblasts (**Figure 3C**). Furthermore, using an antibody against NF-κB p100/52 showed a similar result, indicating that IκBζ forms a complex with p65 and p100/52 (**Figure 3D**).

**Figure 3.**
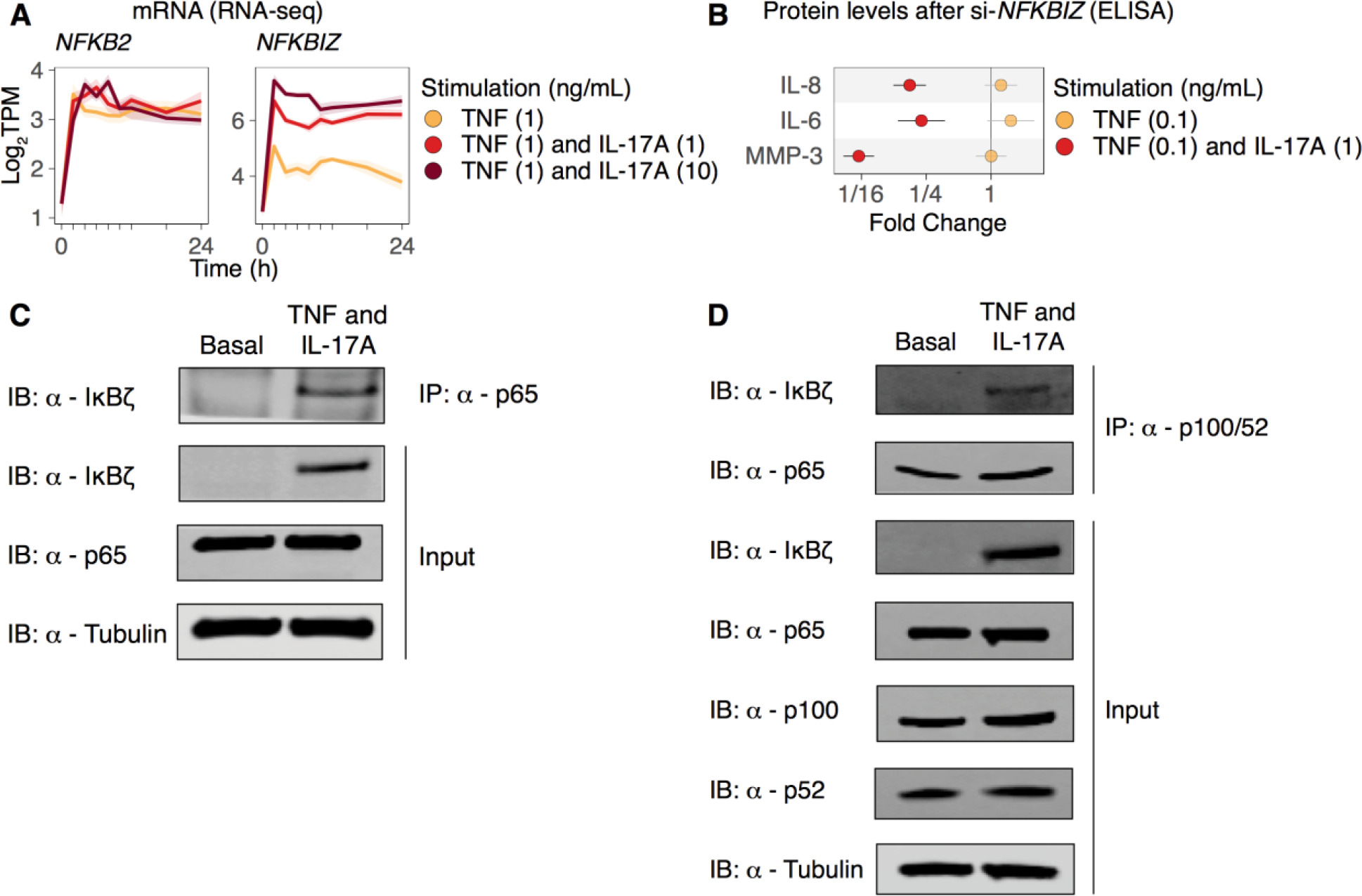
IκBζ binds NF-κB p65 and p100/52, and mediates the response to TNF and IL-17A. **(A)** Gene expression of *NFKB2* (p100) and *NFKBIZ* (IκBζ) over time and IL-17A dosage in **RNA-seq Data 1**. **(B)** Silencing *NFKBIZ* reduces protein levels of MMP-3, IL-6, and IL-8 after costimulation with TNF (0.1 ng/mL) and IL-17A (1 ng/mL), but not after stimulation with TNF alone. **(C)** After 4 hours of costimulation with TNF (1 ng/mL) and IL-17A (1 ng/mL), IκBζ is detected in a pull down of NF-κB p65. **(D)** After 4 hours of costimulation with TNF (1 ng/mL) and IL-17A (1 ng/mL), IκBζ is detected in a pull down of NF-κB p100/52.

### CUX1 and IκBζ mediate neutrophil and monocyte recruitment

Neutrophils are the most abundant cells in the synovial fluid of inflamed joints in RA (Hollingsworth et al., 1967; Zvaifler, 1973), and abundance of synovial macrophages correlates with radiologic score of joint damage (Mulherin et al., 1996) and disease activity (Haringman et al., 2005). So, we performed *in vitro* assays to test which mediators are required for recruitment of neutrophils and monocytes. Briefly, we stimulated synovial fibroblast cell lines with TNF and IL-17A for 24 hours after silencing *CUX1*, *LIFR*, *STAT3*, *STAT4*, or *NFKBIZ*. Then, we tested the supernatants in a transwell migration experiment (**Figure 4A**, **Methods**).

**Figure 4.**
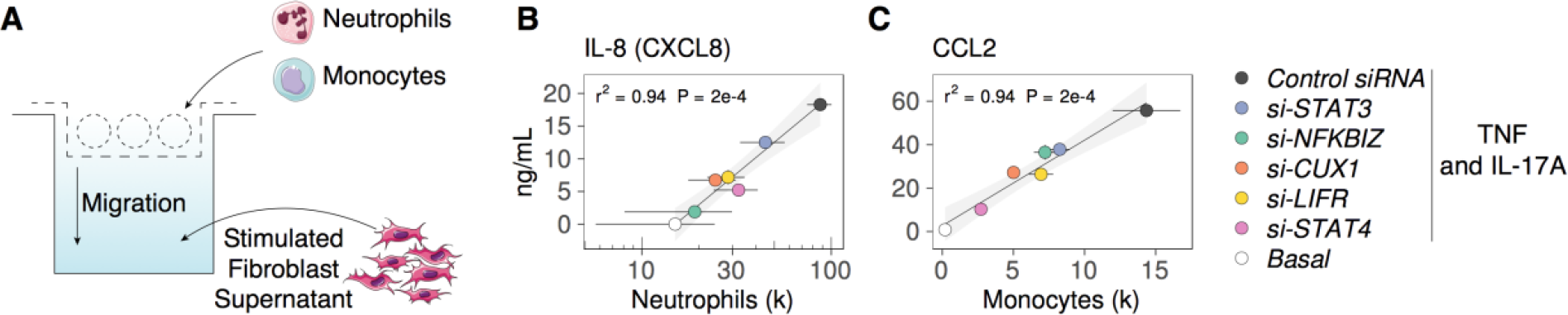
CUX1 and IκBζ mediate neutrophil and monocyte recruitment. **(A)** We tested migration and protein levels in supernatant from synovial fibroblasts before stimulation (Basal) and 24 hours after stimulation with TNF and IL-17A. **(B)** Silencing *CUX1* or *NFKBIZ* (IκBζ) reduces neutrophil recruitment in an *in vitro* migration assay. Neutrophil migration is correlated with protein level of IL-8 in the supernatant measured by ELISA. **(C)** Silencing *CUX1* or *STAT4* reduces monocyte recruitment in an *in vitro* migration assay. Monocyte migration is correlated with protein level of CCL2 in the supernatant measured by ELISA.

After silencing *CUX1* or *LIFR*, the supernatant from synovial fibroblasts was less effective at inducing migration of neutrophils (t-test P = 0.05 and 0.06, respectively) (**Figure 4B**). The number of migrated cells is proportional to protein level of IL-8 (CXCL8) in the supernatant, and silencing *NFKBIZ* had the greatest effect. While monocyte migration was reduced after silencing of *CUX1* or *NFKBIZ* (t-test P = 0.04 and 0.06, respectively) (**Figure 4C**), silencing *STAT4* had the greatest effect on monocyte migration and CCL2 protein level. We conclude that CUX1 and IκBζ mediate recruitment of neutrophils and monocytes.

### Genes induced by TNF and IL-17A are highly expressed in RA synovial tissues

To test if genes induced by TNF or TNF and IL-17A are highly expressed *in vivo*, we examined public RNA-seq data from two large-scale transcriptomics studies that include inflamed RA synovial tissues (Guo et al., 2017; Zhang et al., 2018). We focused on the 51 genes in Categories C1 or C2 and tested for differential gene expression in each study. First, we compared 57 synovial tissues (all cell types) from early RA versus 28 normal tissues (Guo et al., 2017) (**Figure 5A**). 14 of 23 genes in Category C1 and 20 of 25 genes in Category C2 were higher in early RA (FDR < 0.05). Since this study gave us a glimpse of the transcriptomic profile of the whole tissue, we next looked only at the contribution from fibroblasts. In the second study, sorted CD45^-^PDPN^+^ fibroblasts were isolated from synovial tissues, and here we compare 16 samples from inflamed RA tissues versus 29 from non-inflamed tissues (lower abundance of infiltrating leukocytes) (Zhang et al., 2018) (**Figure 5B**). 2 of 26 genes in Category C1 (*LYPD3* and *NR4A2*) and 9 of 25 genes in Category C2 (including *CXCL1*, *CXCL2*, *CXCL3, NFKBIZ*) were higher in fibroblasts from inflamed RA than non-inflamed arthritis tissues (FDR < 0.05). This suggests that fibroblasts may be an important source of chemokines encoded by *CXCL1*, *CXCL2*, *CXCL3, CXCL6, CXCL8, IL6, CCL2, CCL8.* This highlights the relevance of CUX1 and IκBζ mediated transcription in inflamed human tissue.

**Figure 5.**
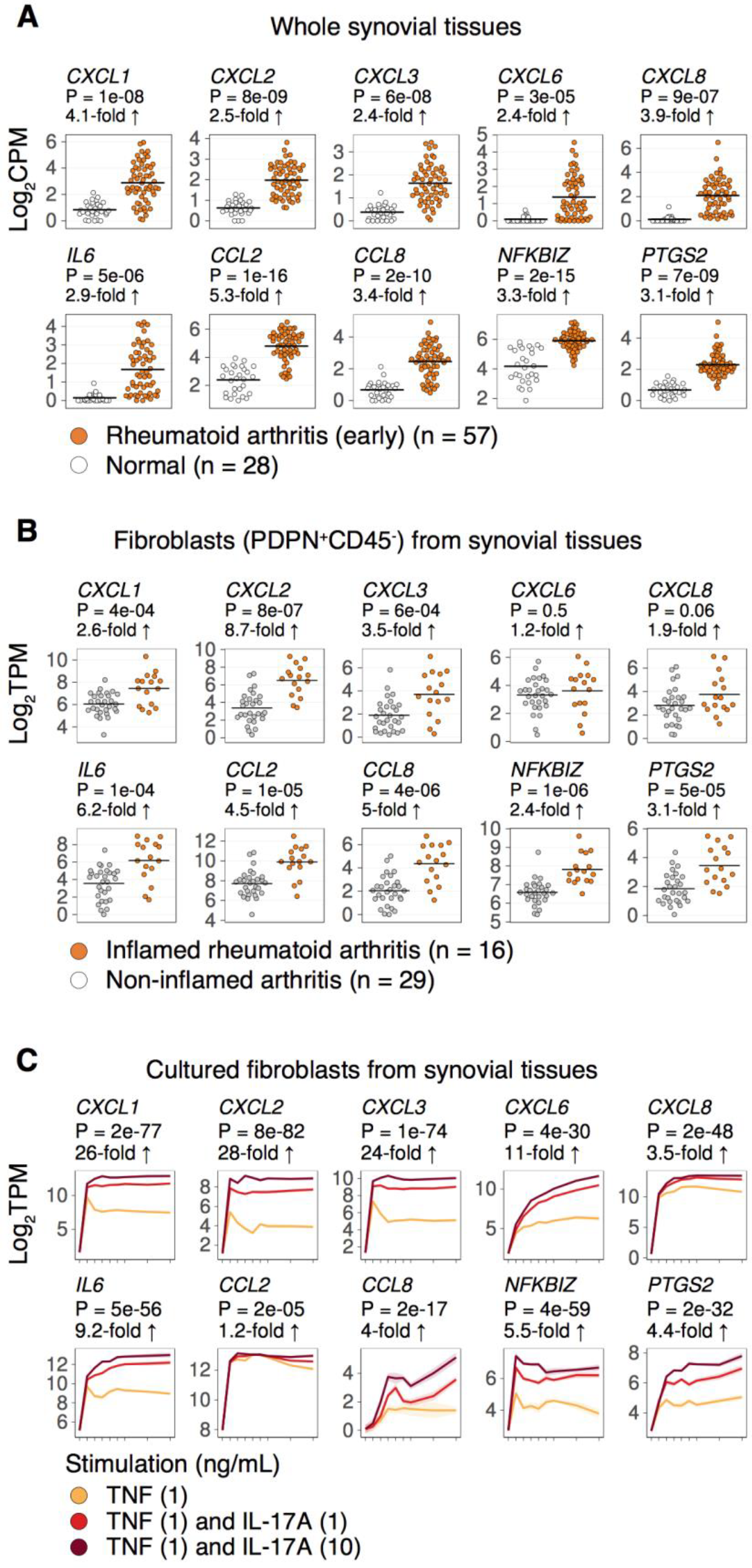
Genes induced by TNF and IL-17A are highly expressed in RA synovial tissues. Transcript abundances from **(A)** RNA-seq data for whole synovial tissues or **(B)** RNA-seq data for sorted synovial fibroblasts (CD45^-^PDPN^+^) from synovial tissues. Each dot represents one sample. Fold-changes and t-test p-values. **(C)** Transcript abundances from **RNA-seq Data 1**, showing abundance over time and IL-17A dose. Fold-changes and t-test p-values for TNF (1 ng/mL) and IL-17A (10 ng/mL) versus TNF (1 ng/mL).

## Discussion

We provide a genome-wide view of transcriptional responses over time to the combination of TNF and IL-17A and to TNF alone. Since IL-17A by itself has weak transcriptional effects, we used time series transcriptomics after stimulation with TNF and three dosages of IL-17A (0, 1, 10 ng/mL) across seven human donor derived cell lines. Multiple dosages of IL-17A allowed us to confidently identify genes with transcriptional abundance proportional to IL-17A dosage. We defined categories of genes by looking at their transcriptional responses to TNF alone and costimulation with TNF and IL-17A. For example, many genes known to be transcriptionally induced by interferon alpha and interferon gamma are induced by TNF alone and repressed by the addition of IL-17A. By querying public RNA-seq data, we also showed that many genes induced by TNF and IL-17A are more highly expressed in inflamed RA synovial tissues, and in particular in synovial fibroblasts from RA joint tissues (**Figure 5**).

TNF is known to induce fibroblasts to produce many inflammatory cytokines, and IL-17A has been shown to synergize with TNF to further induce key cytokines such as IL-6 (Chabaud et al., 2001; Fossiez et al., 1996; Ruddy et al., 2004; Shen et al., 2005), neutrophil attracting chemokines (Ermann et al., 2014; Zrioual et al., 2009), and matrix metalloproteinases (Koshy et al., 2002; Moran et al., 2009). Two transcription factors, NFκB and C/EBP, control synergistic production of IL-6 (Moran et al., 2009; Ruddy et al., 2004) in response to the combination of TNF and IL-17A.

Among the list of IL-17A dose-responsive genes, we saw known IL-17A mediators with dose-responsive transcription such as *NFKBIZ* (Muromoto et al., 2016) and *ZC3H12A* (Koga et al., 2011). We used motif enrichment analysis to discover a unique CUX1-NF-κB motif and predicted that CUX1 mediates transcription of CXC chemokines and that CUX1 might form a complex with NFKB. We validated this prediction by showing that silencing CUX1 with siRNA represses transcription of the genes with the CUX1-NF-κB motif (*CXCL1, CXCL2, CXCL3*). Moreover, CUX1 binds NF-κB p65 after costimulation with TNF and IL-17A. We found increased CUX1 binding to the promoters of *CXCL1, CXCL2, CXCL3* after stimulation with TNF and IL-17A, suggesting that a CUX1 NF-KB complex directly regulates transcription of these chemokines. Our data suggests that CUX1 mediated transcription is independent of LIFR, STAT3, STAT4, and ELF3, because silencing these mediators did not repress transcription of *CXCL1, CXCL2,* and *CXCL3*.

CUX1 is a homeobox transcription factor with ubiquitous expression across mammals and essential roles in development and homeostasis (Hulea and Nepveu, 2012), but its role in inflammation is not well understood. Previously, in tumor associated macrophages, CUX1 was found to bind NF-κB p65 and suppress chemokines *CXCL10* and *CCL5* transcription (Kühnemuth et al., 2013). Here, in fibroblasts, we find that the CUX1 NF-κB p65 complex induces chemokine transcription, suggesting cell type-specific regulation. In future studies, we might expect to find that additional homeobox transcription factors work with NF-κB to regulate inflammation, with different cell-type specific effects.

Previously, IκBζ, a unique member of the IκB family with a distinct role as a coactivator of NF-κB, was shown to be induced in macrophages after stimulation with a number of different toll-like receptor (TLR) ligands and IL-1β, but not after stimulation with TNF alone (Yamamoto et al., 2004). IκBζ regulates transcription of a specific set of genes such as *Csf3* in mouse embryonic fibroblasts (Yamazaki et al., 2008). Here, we found that IκBζ is one of the critical mediators for the synergistic response to TNF and IL-17A in fibroblasts. By silencing *NFKBIZ*, the gene encoding IκBζ, production of IL6 and MMP3 was reduced after costimulation with TNF and IL-17A, but not after stimulation with TNF alone. We also found that synovial fibroblasts require IκBζ for transcription of *CXCL8*, for production of the encoded protein IL-8, and for recruitment of neutrophils *in vitro*. Thus, IκBζ has a critical role in driving inflammation after activation of fibroblasts.

IL-17A is an important cytokine for pathogenesis in autoimmune and inflammatory diseases (Miossec, 2017; Onishi and Gaffen, 2010) such as psoriasis, psoriatic arthritis, ankylosing spondylitis, rheumatoid arthritis (RA) (Lubberts, 2015), inflammatory bowel disease (IBD), and multiple sclerosis (Hu et al., 2011). Two antibodies targeting IL-17A have shown excellent results for treatment of psoriasis: secukinumab is approved for psoriasis, psoriatic arthritis, and ankylosing spondylitis (R.G. et al., 2014), and ixekizumab is approved for psoriasis (Gordon et al., 2016). One might have expected similar results for RA, since IL-17A knockout suppresses arthritis in a collagen-induced arthritis mouse model (Chu et al., 2007), and mouse models have shown IL-17A involvement in all stages of rheumatoid arthritis (Benedetti and Miossec, 2014). However, clinical trials of IL-17A inhibitors with RA patients have shown mixed results (Miossec and Kolls, 2012). There are several possible explanations for this discrepancy. One possibility we might expect is that the heterogeneity in clinical response to IL-17A inhibition could be partially explained by genetic variation in the components of regulatory pathways downstream of TNF and IL-17A.

## Acknowledgments

This work is supported in part by funding from the National Institutes of Health (U01GM092691, UH2AR067677, U19AI111224 (SR)), the Doris Duke Charitable Foundation Grant #2013097, the Ruth L. Kirschstein National Research Service Award (F31AR070582) from the National Institute of Arthritis and Musculoskeletal and Skin Diseases (KS). We thank Joerg Ermann for valuable comments.

## Author Contributions

The manuscript was written by KS with help from SR. All data analysis was done by KS. ELISA experiments were done by EHN and HNN. **Microarray Data 1 and 2** experiments were designed and performed by EHN, GFMW, FM, and HNN. **RNA-seq Data 1 and 2** were designed by KS, HNN, SR, and performed by HNN and GFMW. ChIP, QCPR, western blot, and migration data were collected by HNN, DPS, and MFG. MBB and SR supervised the study.

---

**Figure S1.**
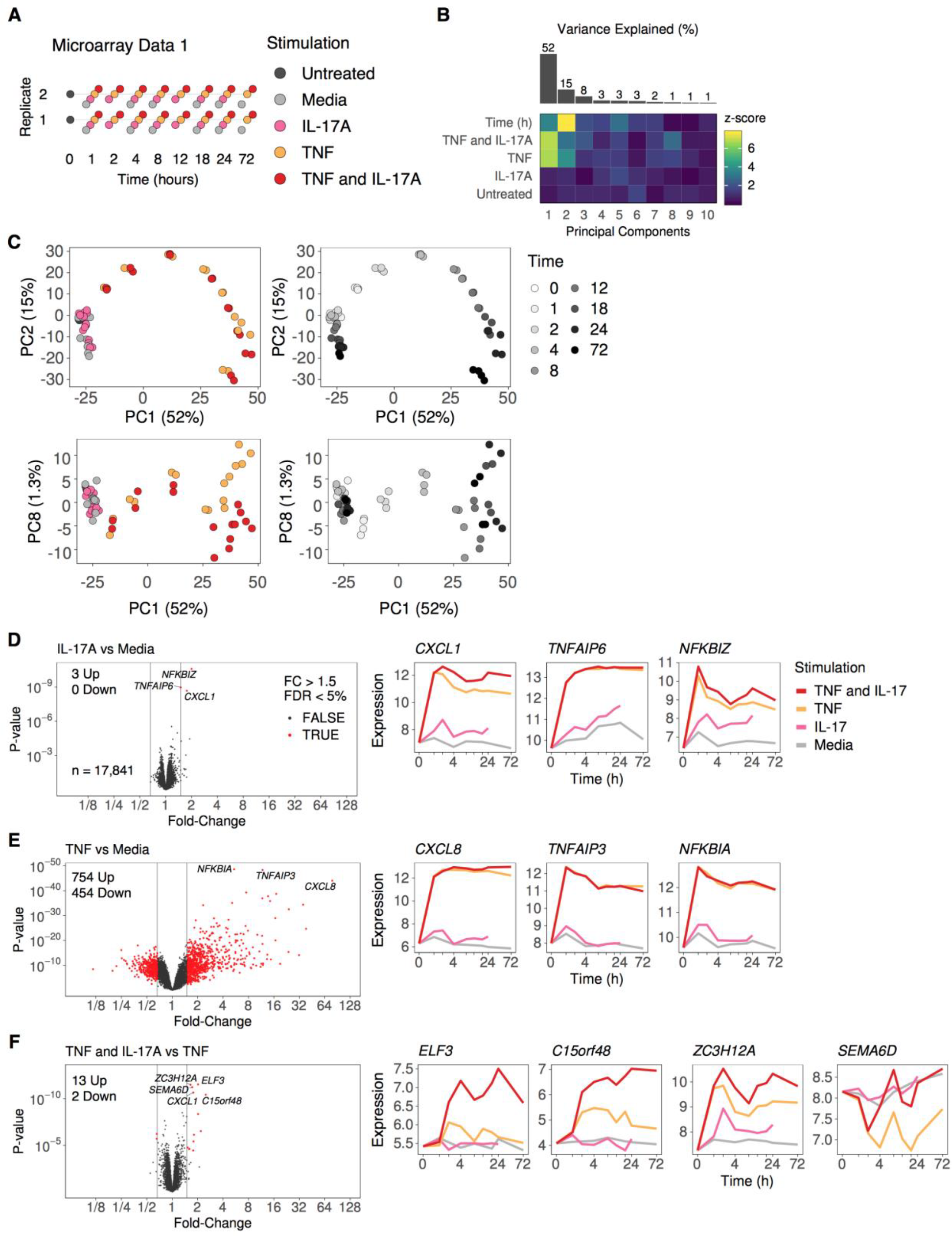
Related to Figure 1 and Table S1. IL-17A requires TNF to induce gene expression response in fibroblasts. **(A)** Experimental design for **Microarray Data 1** (n = 60): 9 time points, 2 technical replicates, 5 different stimulations (Untreated, Media, IL-17A (1 ng/mL), TNF (1 ng/mL), TNF and IL-17A). We used the GeneChip™ Human Gene 1.0 ST Array ("affy_hugene_1_0_st_v1" on Bioconductor). **(B)** Summary of principal components analysis (PCA) on 17,841 genes. We see 52, 15, and 8 percent of variation is captured by the first 3 principal components (PCs). We fit a linear model to each PC (*PC ~ Time + Treatment*) where *Time* is a numeric variable and *Treatment* is a factor. Then we transformed the term p-values to normal z-scores and displayed them in a heatmap. We see that PCs 1 and 2 have strongest association with time, and PCs 1 and 8 have the strongest association with TNF and IL-17A costimulation. **(C)** Samples projected onto PC1 and PC2 (top) or PC1 and PC8 (bottom), colored by stimulation (left) or time (right). Along the first two principal components, stimulation with TNF and IL-17A is similar to TNF alone. Also, stimulation with IL-17A alone is similar to no stimulation at all (Media). **(D, E, F)** Volcano plots and expression profiles of selected genes. We set a threshold at 0.05 FDR and 1.5 fold-change. At this threshold, 3 genes are induced by IL-17A versus media **(D)**, 754 by TNF versus media **(E)**, and 13 by TNF and IL-17A versus TNF **(F)**. 454 genes are repressed by TNF, and 2 genes repressed by TNF and IL-17A versus TNF. See the full list in **Table S1**. Adjacent panels show gene expression levels over time, colored by stimulation. Lines represent means of 2 observations for each stimulation. The y-axis represents microarray expression intensity on the Log_2_ scale.

**Figure S2.**
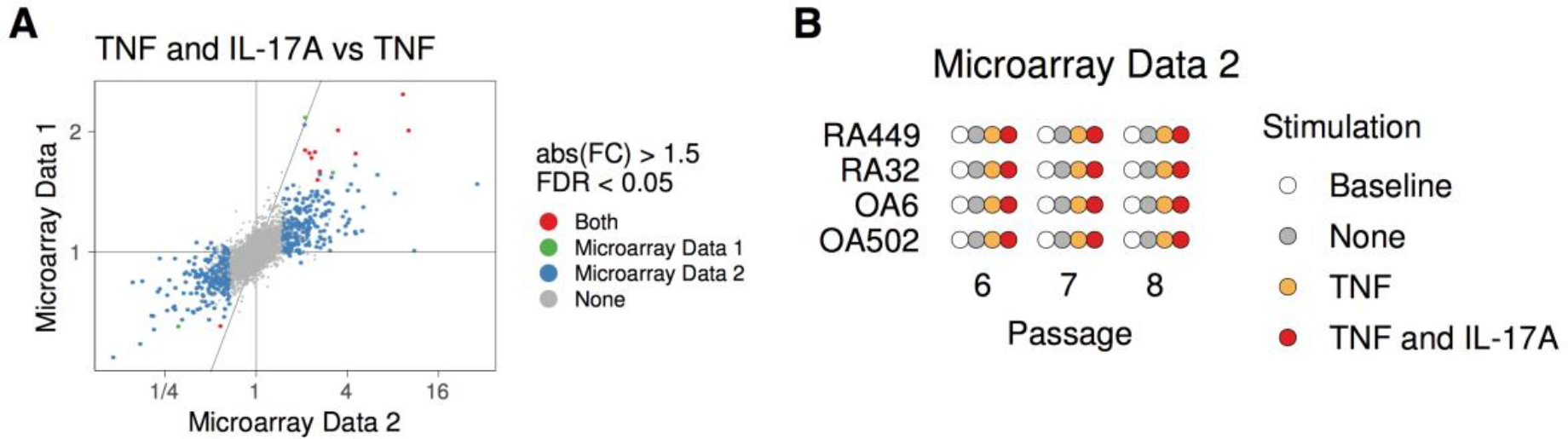
Related to Figure 1 and Tables S1 and S2. Synergy between TNF and IL-17A is reproducible across independent experiments, laboratories, technologies, and cell lines. **(A)** We tested differential expression in **Microarray Data 1** and **Microarray Data 2**, and found that the log2 fold-change is correlated (Pearson correlation = 0.67, p < 2 × 10^-16^). We found 11 genes with significant differential expression in both datasets for the contrast of TNF and IL-17A versus TNF: *ABCA8, AGPAT4, C15orf48, CCL8, CFAP69, CNKSR3, CXCL1, CXCL2, EDNRB, ELF3, SEMA6D*. **(B)** Experimental design for **Microarray Data 2** (n = 48): 1 time point (24 hours), 3 passages, 4 cell lines, 4 different stimulations (Baseline, None, TNF, TNF and IL-17A). We used the GeneChip™ Human Genome U133 Plus 2.0 Array ("affy_hg_u133_plus_2" on Bioconductor).

**Figure S3.**
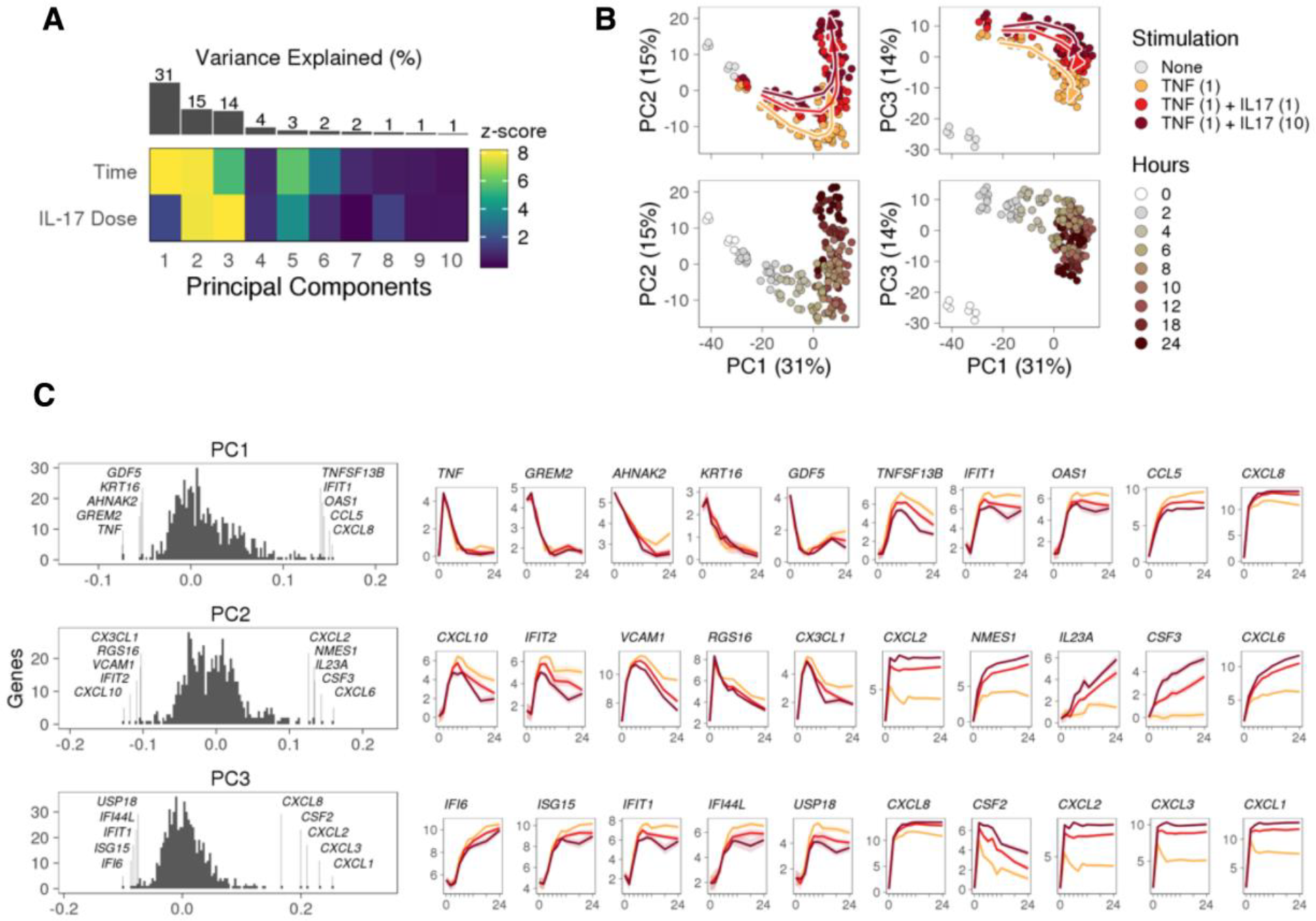
Related to Figure 1. Principal Components Analysis (PCA) of RNA-seq Data 1. **(A)** Summary of PCA results on 636 genes. We see 31, 15, and 14 percent of variation is captured by the first 3 principal components (PCs). We fit a linear model to each PC (*PC ~ Time + Dose*) where *Time* is a numeric variable and *Dose* is a numeric variable. Then we transformed the term p-values to normal z-scores and displayed them in a heatmap. We see that PCs 1 and 2 have strongest association with time and PC3 has the strongest association with IL-17A dose. **(B)** Samples projected onto PC1 and PC2 (left) or PC1 and PC3 (right), colored by stimulation (top) or time (bottom). **(C)** Loading values for 636 genes, highlighting the top 5 genes with lowest and top 5 genes with highest values. We see that each principal component captures different patterns of variation over time and response to IL-17A dose.

**Figure S4.**
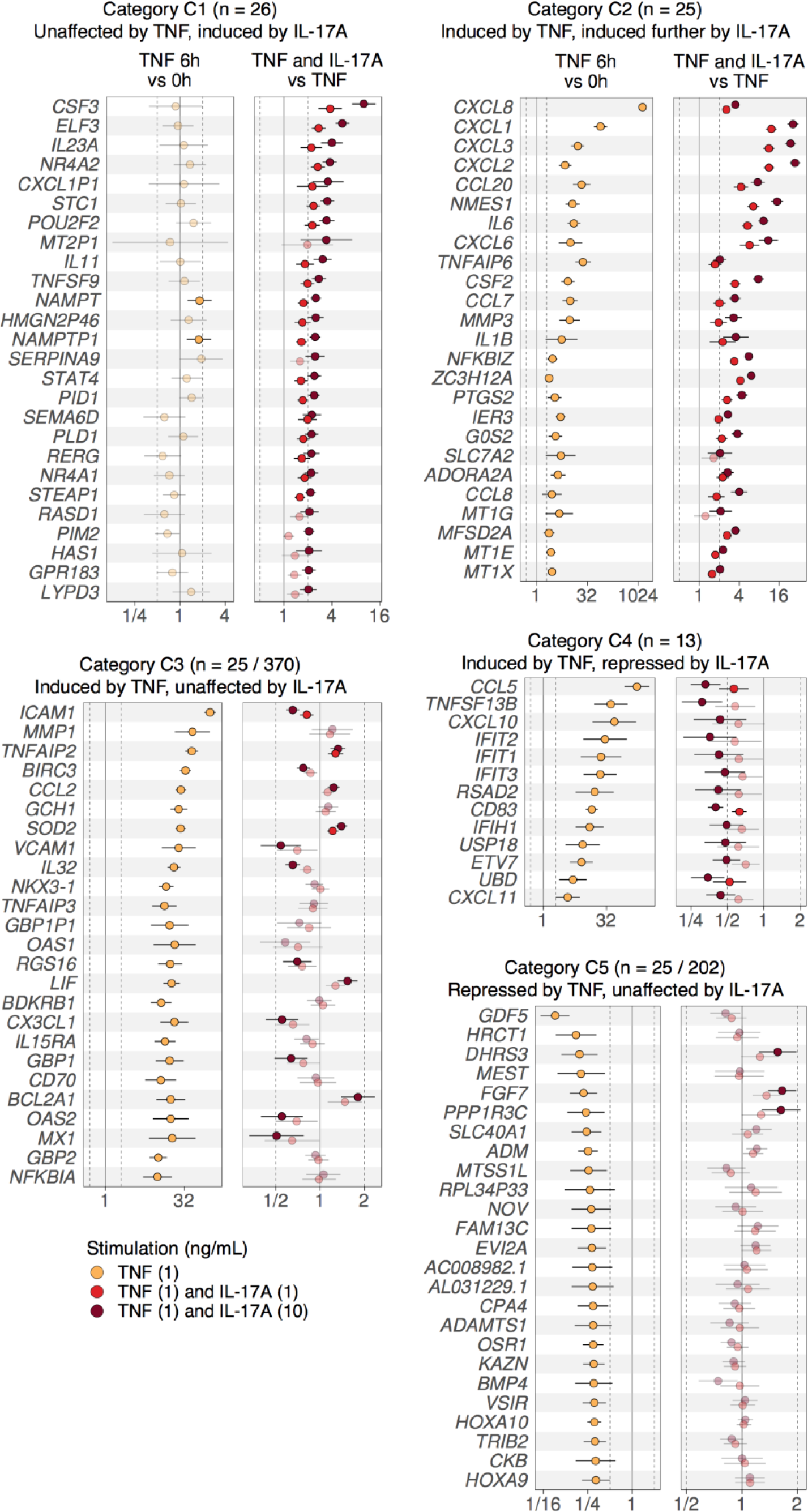
Related to Figure 1 and Table S3. Response to TNF alone or the combination of TNF and IL-17A. Differential gene expression effect sizes and 95% confidence intervals for genes in each of the 5 Categories (C1, C2, C3, C4, C5). Three contrasts are shown: TNF at 6h versus 0h, TNF and IL-17A (1 ng/mL) vs TNF, and TNF and IL-17A (10 ng/mL) vs TNF. Faded color indicates FDR > 0.05.

**Figure S5.**
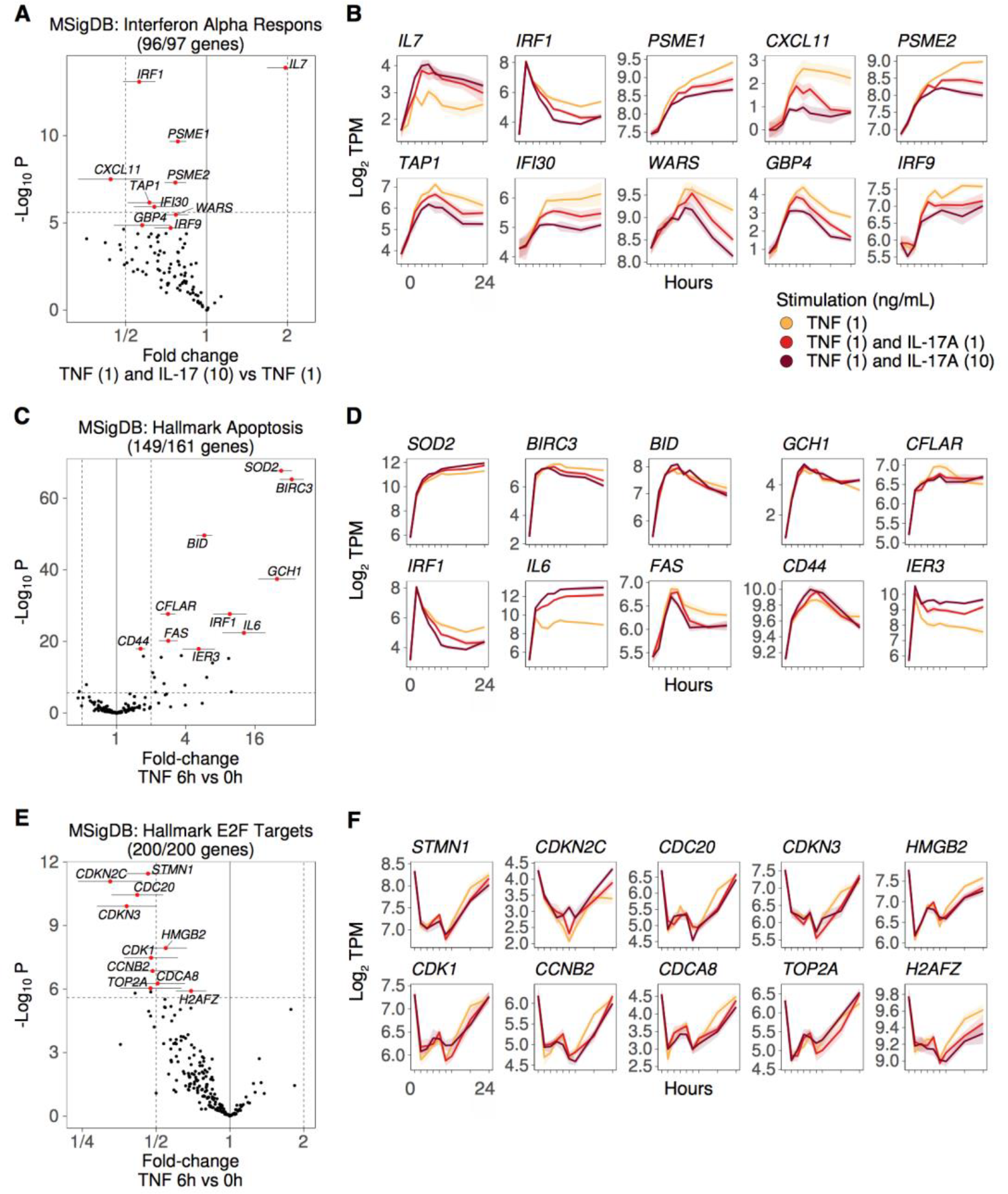
Related to Figure 1. Genes induced by TNF in the MSigDB Hallmark gene set called "Interferon Alpha Response" are repressed by IL-17A. **(A,C,E)** Volcano plots, showing fold change estimates and p-values for the contrast of TNF and IL-17A (10 ng/mL) versus TNF. The top 10 genes with lowest p-values are highlighted in red. The titles indicate the number of genes measured in **RNA-seq Data 1** and the total number of genes in the gene set, e.g. 96 of the 97 genes in the MSigDB Hallmark gene set called "Interferon Alpha Response" are measured in **RNA-seq Data 1**. **(B,D,F)** Expression over time, shown as mean +/-SEM of the Log2 transcripts per million (TPM).

**Figure S6.**
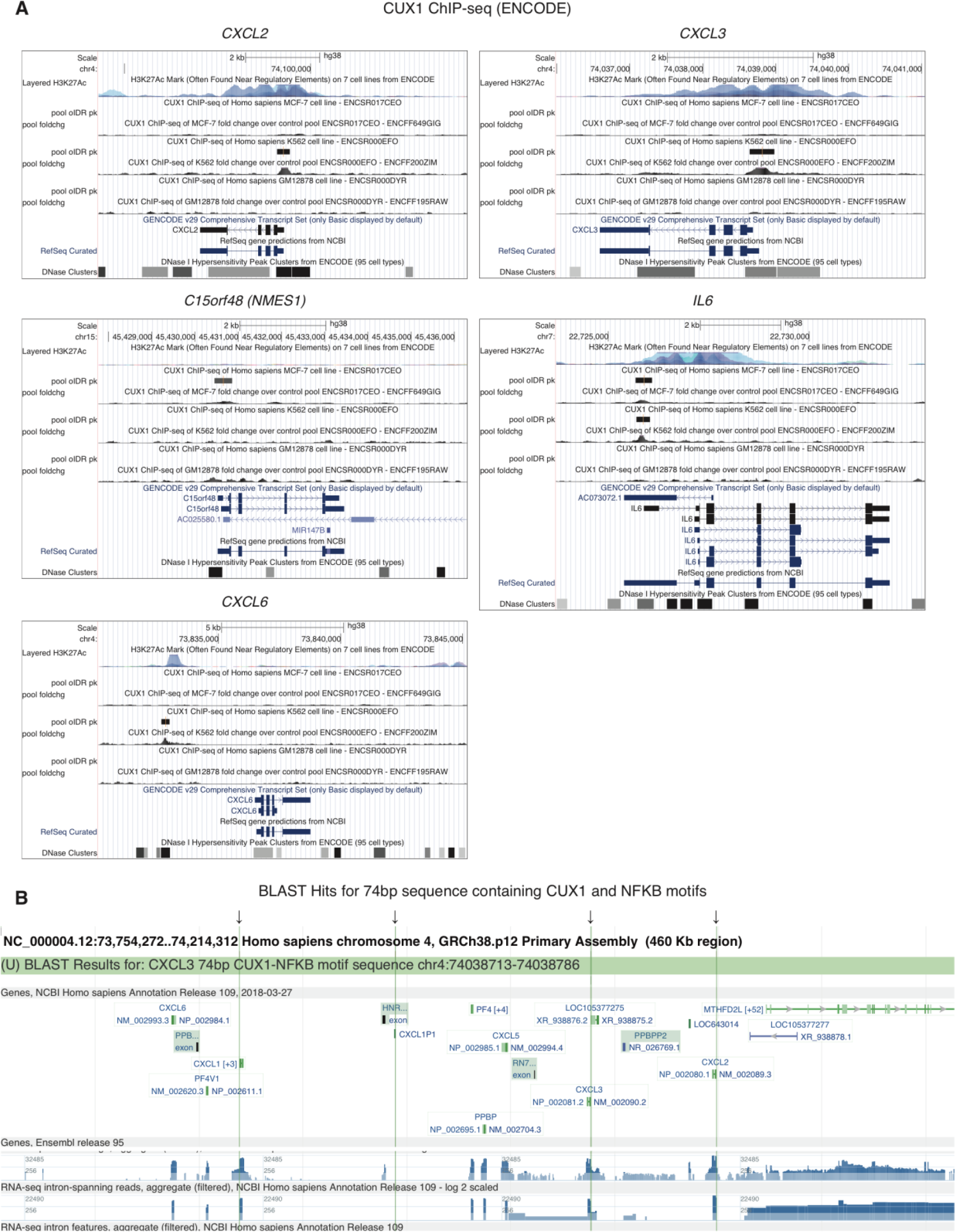
Related to Figure 2. ENCODE CUX1 ChIP-seq peaks in promoters of *CXCL2*, *CXCL3*, *C15orf48* (*NMES1*), *IL6*, and near *CXCL6*. (**A**) We used UCSC Genome Browser to visualize CUX1 ChIP-seq data (https://www.encodeproject.org/targets/CUX1-human/) in three cell lines: MCF-7, K562, GM12878. ChIP-seq tracks show fold change over control and optimal IDR peak calls. We also show H3K27Ac (7 cell lines from ENCODE) and DNase 1 hypersensitivity peak clusters from ENCODE (95 cell types). (**B**) A 74 bp sequence with motifs for NFKB1, CUX1, and STAT3 is only present in the promoters of *CXCL1, CXCL2, CXCL3,* and *CXCL1P1*. We used the following human genomic sequence of 74 nucleotides (hg38 chr4:74038713-74038786 strand=+) as a query with NCBI BLAST: GGATCGATCTGGAGCTCCGGGAATTTCCCTGGCCCGGCCGCTCCGGGCTTTCCAGTCTCAACCATGCATAAAAA We queried against “Human genomic plus transcript (Human G + T)” and found 7 hits. 3 hits were on transcripts (*CXCL3* mRNA, *CXCL2* mRNA, and uncharacterized LOC105377275). 4 hits were on genomic sequences, shown in this figure to be located at the promoters of *CXCL1, CXCL2, CXCL3,* and pseudogene *CXCL1P1*.

**Figure S7.**
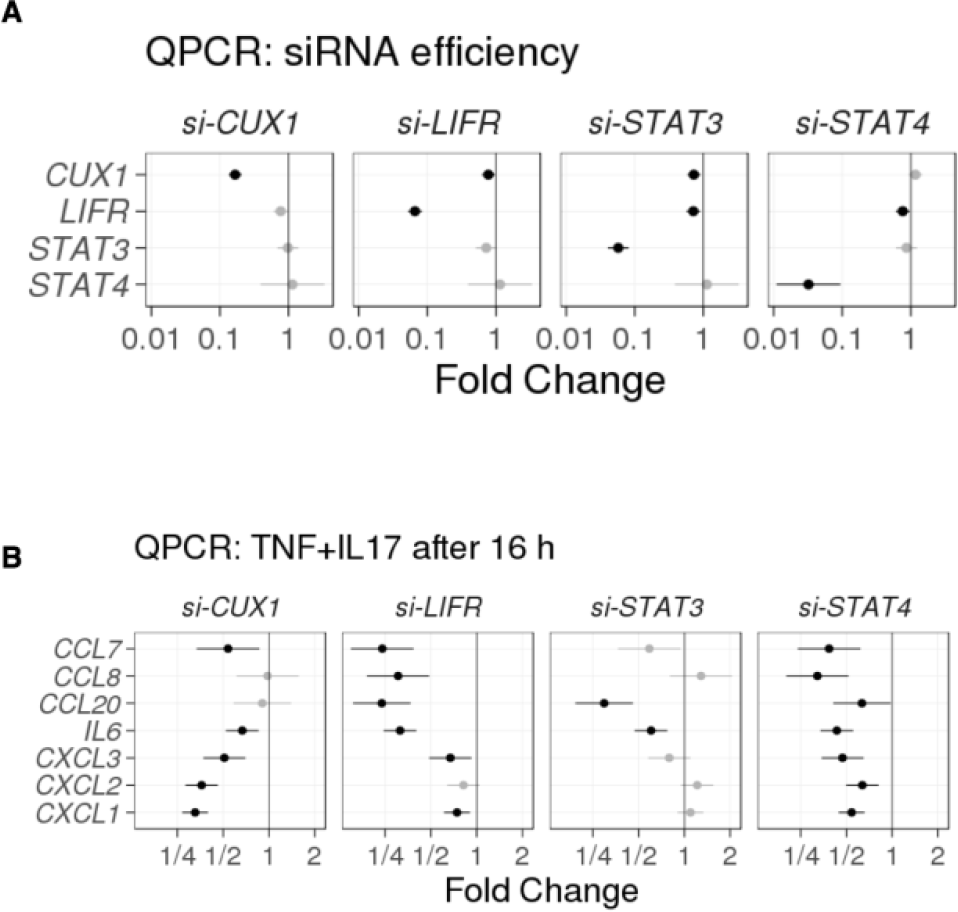
Related to Figure 2. Efficiency of silencing each transcription factor with siRNA, and effects of transcription factor silencing on selected genes. (**A**) *CUX1*, *LIFR*, *STAT3*, and *STAT4* were effectively silenced by siRNA. We cultured 4 RA synovial fibroblast cell lines and measured silencing efficiency in each one. For *CUX1*, *LIFR* and *STAT3* siRNA, the silencing efficiency was measured at basal conditions. For *STAT4* siRNA, it was quantified 16 hours after costimulation with TNF and IL-17A. (**B**) qPCR analysis of the cDNA libraries prepared for RNA-seq. We show results for *CCL7*, *CCL8*, *CCL20*, *IL6*, *CXCL3*, *CXCL2*, *CXCL1*. After incubating 4 cell lines with siRNAs for *CUX1, LIFR, STAT3,* or *STAT4*, we measured the selected genes with qPCR relative to *GAPDH*. Transcription of *CXCL1, CXCL2,* and *CXCL3* is repressed after silencing *CUX1*, but not as much after silencing *LIFR, STAT3,* or *STAT4.* Transcription of *CCL7, CCL8, CCL20,* and *IL6* is repressed after silencing *LIFR*, but not as much after silencing *CUX1*. Dots represent mean fold change relative to control siRNA, and error bars represent 95% confidence intervals. Black color indicates FDR < 0.05.

## Supplemental Tables

**Table S1. Full analysis of Microarray Data 1.**

We calculated fold-changes, 95% confidence intervals, p-values, and q-values for each gene: effect of IL-17A versus media, TNF versus media, and TNF and IL-17A versus TNF.

**Table S2. Full analysis of Microarray Data 2.**

We calculated fold-changes, 95% confidence intervals, p-values, and q-values for each gene: effect of TNF versus media, TNF and IL-17A versus media, and TNF and IL-17A versus TNF.

**Table S3. Full analysis of RNA-seq Data 1.**

We calculated fold-changes, 95% confidence intervals, p-values, and q-values for each gene: effect of TNF at each time point versus 0h, TNF and IL-17A (1 ng/mL) versus TNF, and TNF and IL-17A (10 ng/mL) versus TNF. This table also includes the category for each gene, as defined in Figure 1D.

**Table S4. Full analysis of RNA-seq Data 2.**

We calculated average expression, 95% confidence intervals, p-values, and q-values for each gene: average expression for control siRNA at 0, 1, 6, 16h after stimulation with TNF and IL-17A (0 or 1 ng/mL IL-17A).

**Table S5. Sequences of primers used in qPCR and ChIP assays.**

We used the provided sequences for qPCR and ChIP experiments assaying IL-6, GAPDH, STAT3, STAT4, CUX1, LIFR, CXCL1, CXCL2, CXCL3, LIF, IL-8, CCL20.

## Methods

### Cell Lines and Reagents

Human synovial fibroblasts were isolated as previously described (Kiener et al., 2006) from tissues discarded after synovectomy or joint replacement surgery. Each line was derived from a unique donor and used experimentally between passages 5 and 8. Cells were maintained in Dulbecco’s modified Eagle’s medium (DMEM, Gibco) supplemented with 10% fetal bovine serum (FBS; Gemini), 2 mM L-glutamine, 50 µM 2-mercaptoethanol, antibiotics (penicillin and streptomycin), and essential and nonessential amino acids (Life Technologies). The following antibodies were used: anti-CUX1 (EMD Millipore); anti-p100/p52, anti-IκBζ (Cell Signaling Technology); anti-p65 (Santa-Cruz); anti-β-tubulin (Sigma); anti-CD45, anti-CD11b, anti-CD66b, (BioLegend); anti-CD14 (eBioscience). Other reagents were purchased from the following vendors: IL-6, IL-8, CCL2 and MMP3 ELISA kits, TNF, IL-17A (R&D Systems). All siRNAs (Silencer Select) were purchased from Life Technologies.

### Immunoprecipitation

Total cell lysates were collected by washing cells once with cold PBS followed by addition of lysate buffer (50mM HEPES pH 7.5, 5% glycerol, 100 mM NaCl, 0.25% TritonX-100, supplemented with a protease inhibitor cocktail (Roche) and phosphatase inhibitors (sodium orthovanadate, sodium fluoride, and beta-glycerol phosphate). Cells were lysed for 30 minutes on ice followed by centrifugation at 15000rpm for 15 minutes at 4°C. Immunoprecipitation was done overnight at 4°C using Protein A resins together with a rabbit antibody against the relevant target.

### Quantitative Real-Time PCR

mRNA samples were extracted from cells using an RNeasy Micro Kit (Qiagen). cDNA synthesis was carried out using the QuantiTect Reverse Transcription Kit (Qiagen). qPCR reactions were performed in duplicate using the Brilliant III Ultra-Fast SYBR reagent (Agilent). qPCR primers are listed in **Table S5**. Relative transcription level was calculated by using the ΔΔCt method with GAPDH as the normalization control. Fold induction was calculated by dividing the normalized mRNA at a certain time point with that at time 0hr. Fold change is the ratio of the normalized mRNA from cells expressing a specific siRNA vs. Ctrl siRNA.

### Chromatin Immunoprecipitation Assays

Fibroblasts were stimulated with TNF (1 ng/mL) + IL-17A (1 ng/mL) for different durations of time as indicated followed by fixation with formaldehyde (1% final concentration) for 10 minutes. Cells were lysed in swelling buffer for 15 minutes (25mM HEPES pH 7.8, 1.5 mM MgCl_2_, 10mM KCl, 0.1% NP-40, 1mM DTT, protease inhibitor cocktail) then in sonication buffer (50mM HEPES pH 7.8, 140mM NaCl, 1mM EDTA, 0.1% SDS, 1% TritonX-100 supplemented with protease inhibitor cocktail). Samples were sonicated for 2 minutes with 20-second pulse intervals and 1 minute off at each interval at 4°C using a Qsonica sonicator set at 28% output. Chromatin was immunoprecipitated using Dynabead Protein A resins (Life Technologies) together with a rabbit antibody against the relevant target at 4°C overnight. DNA was purified using the phenol/chloroform precipitation method. The amount of DNA precipitated was quantified using qPCR. Fold recruitment was calculated by dividing the amount of ChIP-ed DNA at indicated time point (normalized with input DNA) with that at 0hr (normalized with input DNA). ChIP primers are listed in **Table S5**.

### Western Blotting

Total cell lysates were collected by washing cells once with cold PBS followed by addition of lysate buffer (50mM HEPES pH 7.5, 10% glycerol, 100 mM NaCl, 0.1% SDS, 1% NP-40, supplemented with protease inhibitors and phosphatase inhibitors sodium orthovanadate, sodium fluoride, and beta-glycerol phosphate). Cells were lysed for 30 minutes on ice followed by centrifugation at 15000rpm for 15 minutes at 4°C. Protein concentration was measured by the microBCA kit (Pierce). Equal amounts of total protein (~20 µg per lane) were separated on an 8% SDS-PAGE gel. Proteins were transferred onto a PVDF membrane and blocked with 5% BSA in PBS and probed with primary antibodies overnight at 4°C, followed by secondary antibodies conjugated with IRDye 680 or 800 (Rockland). Membranes were scanned with a Li-COR Odyssey scanner.

### Leukocyte Migration

Fibroblasts were transfected with a control siRNA, or siRNA against a gene by reverse transfection using the RNAiMax reagent (Life Technologies) on day 1. The following day, cells were switched to 0.5% serum media and after 24 hours they were stimulated with TNF (0.5 ng/mL) + IL-17A (0.5ng/mL) for another 24 hours before supernatants were collected and added to the bottom well of a 24-well plate. Neutrophils (PMNs) purified from blood using the neutrophil isolation kit (Stem Cell Technologies) and peripheral blood mononuclear cells (PBMC) containing monocytes purified from blood using Ficoll isolation technique were then introduced to the top well (Corning, 3µm pores insert) at the concentration of 1 million 1×10^6^ cells per mL for PMNs and 2×10^6^ cells per mL for PBMC. After 75 mins (PMN migration) or 3 hours (monocyte migration), cells from the bottom well were collected and the number of neutrophils, monocytes recruited toward the bottom well were quantified by flow cytometry using (anti-CD66b and anti-CD11b for neutrophils), (anti-CD14 and anti-CD45 for monocytes), and cell numbers were normalized using counting beads (Spherotech).

### RNA-seq Expression Profiling

Fibroblasts were plated on day 1 at 50,000 cells per well in 24-well plates in 10% FBS containing media. Cells were serum-starved on day 2 by changing to 1% FBS-containing media. Cells were either left unstimulated or stimulated with TNF (1 ng/mL) or TNF (1 ng/mL) + IL-17A (1 ng/mL) for various durations of time. Samples were collected and RNA was extracted using the RNeasy Micro Kit (Qiagen). RNA-seq libraries were prepared with the Smart-Seq2 protocol.

### siRNA Silencing

Fibroblasts were transfected with an siRNA by reverse transfection at 30 nM using the RNAi Max reagent (Life Technologies) in 10% FBS containing media. Cells were then switched to serum-starving media containing 1% FBS on day 2. Cells were stimulated as indicated on day 3. Efficiency of siRNA silencing was assessed by qPCR (**Figure S7**).

### Data Visualization

All of the figures were created with R (https://www.R-project.org) and many supporting packages, including: ggplot2, ggrepel, patchwork, RColorBrewer, seriation, viridis.

### Microarray Data Analysis

We normalized the data from 60 microarrays with the robust multi-array averaging (RMA) algorithm (Irizarry et al., 2003) from the **oligo** R package (Carvalho and Irizarry, 2010), downloaded from Bioconductor (Huber et al., 2015). To assess quality of the microarray data, we looked at the Normalized Unscaled Standard-Errors (NUSE) and Relative Log-Expression (RLE), and decided to keep all microarrays in our analysis.

We used the **BioMart** R package (Durinck et al., 2005, 2009) (from Bioconductor) to download gene names and identifiers for all of the oligonucleotide probe identifiers that match the array name "affy_hugene_1_0_st_v1". We averaged all of the probes for each gene.

We assess the quality of expression data for each array as follows. For each array, we computed the median *M* RMA expression value across all genes. For each gene, we computed the percent *P* of microarrays with expression greater than *M*. We defined a set of 7,891 genes with *P* > 90% and call them "common genes". In other words, the common genes had high expression in most microarrays. For each microarray, we computed how many of the common genes had expression greater than *M*. This is the percent of common genes detected for each microarray. All of the 60 microarrays detected 97% or more of the common genes, so we decided to keep all of them in our analysis.

We used the **limma** R package (Ritchie et al., 2015) to fit a linear model to each gene and test for differential gene expression across contrasts of interest. For **Microarray Data 1**, we fit the following model:

> *GeneExpression* ∼ *TimeFactor* + *TreatmentFactor*

*TimeFactor* represents 7 indicator variables, one for each time point (we excluded t = 0). *TreatmentFactor* represents 3 indicator variables, one for each treatment: (1) IL-17A, (2) TNF, and (3) TNF and IL-17A. We illustrate the experimental design in **Figure S1A**.

For **Microarray Data 2**, we fit the following model:

> *GeneExpression* ∼ *TreatmentFactor*

*TreatmentFactor* represents 3 indicator variables, one for each treatment after 24 hours: (1) media, (2) TNF, and (3) TNF and IL-17A. We illustrate the experimental design in **Figure S2B**.

### RNA-seq Data Analysis

We used kallisto (version 0.43.1) (Bray et al., 2016) to quantify transcripts per million (TPM) for all of the transcripts reported in Ensembl release 89 (Aken et al., 2016). To get TPM for each gene, we added the TPM values for all of the transcripts that belong to a gene.

We assessed the quality of each RNA-seq sample with a similar approach to the microarray data analysis. For each gene, we computed the percent *P* of 175 samples with expression greater than 0. We defined a set of 12,087 genes with *P* > 90% and call these the "common genes". These are genes for which expression was detected in most samples. For each sample, we computed how many of the common genes had expression greater than 0. This is the percent of common genes detected for each sample. All of the 175 samples detected 98% or more of the common genes, so we kept all of them in our analysis.

We used the limma R package (Ritchie et al., 2015) to fit a linear model to each gene and test for differential gene expression across contrasts of interest. For RNA-seq Data 1, we fit the following model:

> *GeneExpression* ∼ *TimeFactor* + *DoseFactor*

*TimeFactor* represents 8 indicator variables: one for each time point after stimulation with TNF. *DoseFactor* represents 2 indicator variables: one for TNF and IL-17A (1 ng/mL), and one for TNF and IL-17A (10 ng/mL).

We used the CAMERA algorithm from the LIMMA package to compute gene set enrichment p-values with gene sets from MSigDB (Liberzon et al., 2015).

We used HOMER (Heinz et al., 2010) to find transcription factor binding motifs that are overrepresented in promoter sequences. First, we selected 26 genes with at least 1.5 log2 fold-change (2.83 fold-change) greater expression for 10 versus 1 ng/mL IL-17A at 5% false-discovery rate (FDR). We ran the findMotifs.pl script with the Ensembl gene identifiers for these 26 genes as input to find motifs overrepresented in their promoters relative to the background of all other genes. HOMER accepted 25 of the 26 genes for analysis.

We assessed the power gained in **RNA-seq Data 1** by comparing results from analysis of the full data with 9 time points (0, 2, 4, 6, 8, 10, 12, 18, 24 hours) and results from analysis of a subset of the data with 2 time points (0, 24 hours). In the full analysis and the subset analysis, we fit two linear models (reduced model and full model) and computed the F statistic to test how many genes have a significantly better fit with the full model at FDR < 0.05.

> Reduced model: *GeneExpression* ∼ *TimeFactor*
>
> Full model: *GeneExpression* ∼ *TimeFactor* + *DoseFactor*

We found 253 genes were significantly better fit by the full model than the reduced model when we used 9 time points. In contrast, we found 144 genes when we used 2 time points.

## Supplemental Information

### Data availability

**Microarray Data 1**, **Microarray Data 2**, **RNA-seq Data 1**, and **RNA-seq Data 2** will be available at NCBI GEO upon publication.

## References

Aherne, C.M., McMorrow, J., Kane, D., FitzGerald, O., Mix, K.S., and Murphy, E.P. (2009). Identification of NR4A2 as a transcriptional activator of IL-8 expression in human inflammatory arthritis. Mol. Immunol. 46, 3345–3357.

Aken, B.L., Ayling, S., Barrell, D., Clarke, L., Curwen, V., Fairley, S., Fernandez Banet, J., Billis, K., García Girón, C., Hourlier, T., et al. (2016). The Ensembl gene annotation system. Database 2016.

Benedetti, G., and Miossec, P. (2014). Interleukin 17 contributes to the chronicity of inflammatory diseases such as rheumatoid arthritis. Eur. J. Immunol.

Bottini, N., and Firestein, G.S. (2013). Duality of fibroblast-like synoviocytes in RA: passive responders and imprinted aggressors. Nat. Rev. Rheumatol. 9, 24–33.

Bray, N.L., Pimentel, H., Melsted, P., and Pachter, L. (2016). Near-optimal probabilistic {RNA-seq} quantification. Nat. Biotechnol. 34, 525–527.

Carvalho, B.S., and Irizarry, R.A. (2010). A framework for oligonucleotide microarray preprocessing. Bioinformatics 26, 2363–2367.

Chabaud, M., Durand, J.M., Buchs, N., Fossiez, F., Page, G., Frappart, L., and Miossec, P. (1999). Human interleukin-17: A T cell-derived proinflammatory cytokine produced by the rheumatoid synovium. Arthritis Rheum. 42, 963–970.

Chabaud, M., Page, G., and Miossec, P. (2001). Enhancing effect of IL-1, IL-17, and TNF-alpha on macrophage inflammatory protein-3alpha production in rheumatoid arthritis: regulation by soluble receptors and Th2 cytokines. J. Immunol. 167, 6015–6020.

Chu, C.-Q., Swart, D., Alcorn, D., Tocker, J., and Elkon, K.B. (2007). Interferon-$\gamma$ regulates susceptibility to collagen-induced arthritis through suppression of interleukin-17. Arthritis Rheum. 56, 1145–1151.

Durinck, S., Moreau, Y., Kasprzyk, A., Davis, S., De Moor, B., Brazma, A., and Huber, W. (2005). BioMart and Bioconductor: a powerful link between biological databases and microarray data analysis. Bioinformatics 21, 3439–3440.

Durinck, S., Spellman, P.T., Birney, E., and Huber, W. (2009). Mapping identifiers for the integration of genomic datasets with the R/Bioconductor package biomaRt. Nat. Protoc. 4, 1184–1191.

Elshabrawy, H.A., Volin, M. V, Essani, A.B., Chen, Z., McInnes, I.B., Van Raemdonck, K., Palasiewicz, K., Arami, S., Gonzalez, M., Ashour, H.M., et al. (2018). IL-11 facilitates a novel connection between RA joint fibroblasts and endothelial cells. Angiogenesis 21, 215–228.

Ermann, J., Staton, T., Glickman, J.N., de Waal Malefyt, R., and Glimcher, L.H. (2014). Nod/Ripk2 signaling in dendritic cells activates IL-17A-secreting innate lymphoid cells and drives colitis in T-bet-/-.Rag2-/-(TRUC) mice. Proc. Natl. Acad. Sci. U. S. A. 111, E2559-66.

Fossiez, F., Djossou, O., Chomarat, P., Flores-Romo, L., Ait-Yahia, S., Maat, C., Pin, J.J., Garrone, P., Garcia, E., Saeland, S., et al. (1996). T cell interleukin-17 induces stromal cells to produce proinflammatory and hematopoietic cytokines. J. Exp. Med. 183, 2593–2603.

Gordon, K.B., Blauvelt, A., Papp, K.A., Langley, R.G., Luger, T., Ohtsuki, M., Reich, K., Amato, D., Ball, S.G., Braun, D.K., et al. (2016). Phase 3 Trials of Ixekizumab in Moderate-to-Severe Plaque Psoriasis. N. Engl. J. Med. 375, 345–356.

Guo, Y., Walsh, A.M., Fearon, U., Smith, M.D., Wechalekar, M.D., Yin, X., Cole, S., Orr, C., McGarry, T., Canavan, M., et al. (2017). CD40L-Dependent Pathway Is Active at Various Stages of Rheumatoid Arthritis Disease Progression. J. Immunol. 198, 4490–4501.

Haringman, J.J., Gerlag, D.M., Zwinderman, A.H., Smeets, T.J.M., Kraan, M.C., Baeten, D., McInnes, I.B., Bresnihan, B., and Tak, P.P. (2005). Synovial tissue macrophages: a sensitive biomarker for response to treatment in patients with rheumatoid arthritis. Ann. Rheum. Dis. 64, 834–838.

Heinz, S., Benner, C., Spann, N., Bertolino, E., Lin, Y.C., Laslo, P., Cheng, J.X., Murre, C., Singh, H., and Glass, C.K. (2010). Simple combinations of lineage-determining transcription factors prime cis-regulatory elements required for macrophage and B cell identities. Mol. Cell 38, 576–589.

Hillmer, E.J., Zhang, H., Li, H.S., and Watowich, S.S. (2016). STAT3 signaling in immunity. Cytokine Growth Factor Rev.

Hirota, K., Yoshitomi, H., Hashimoto, M., Maeda, S., Teradaira, S., Sugimoto, N., Yamaguchi, T., Nomura, T., Ito, H., Nakamura, T., et al. (2007). Preferential recruitment of CCR6-expressing Th17 cells to inflamed joints via CCL20 in rheumatoid arthritis and its animal model. J. Exp. Med. 204, 2803–2812.

Hirota, K., Hashimoto, M., Ito, Y., Matsuura, M., Ito, H., Tanaka, M., Watanabe, H., Kondoh, G., Tanaka, A., Yasuda, K., et al. (2018). Autoimmune Th17 Cells Induced Synovial Stromal and Innate Lymphoid Cell Secretion of the Cytokine GM-CSF to Initiate and Augment Autoimmune Arthritis. Immunity.

Hollingsworth, J.W., Siegel, E.R., and Creasey, W.A. (1967). Granulocyte survival in synovial exudate of patients with rheumatoid arthritis and other inflammatory joint diseases. Yale J. Biol. Med. 39, 289–296.

Hu, Y., Shen, F., Crellin, N.K., and others (2011). The IL-17 pathway as a major therapeutic target in autoimmune diseases. Ann. N. Y. Acad. Sci.

Huber, W., Carey, V.J., Gentleman, R., Anders, S., Carlson, M., Carvalho, B.S., Bravo, H.C., Davis, S., Gatto, L., Girke, T., et al. (2015). Orchestrating high-throughput genomic analysis with Bioconductor. Nat. Methods 12, 115–121.

Hulea, L., and Nepveu, A. (2012). CUX1 transcription factors: from biochemical activities and cell-based assays to mouse models and human diseases. Gene 497, 18–26.

Irizarry, R.A., Hobbs, B., Collin, F., Beazer-Barclay, Y.D., Antonellis, K.J., Scherf, U., and Speed, T.P. (2003). Exploration, normalization, and summaries of high density oligonucleotide array probe level data. Biostatistics 4, 249–264.

Khan, A., Fornes, O., Stigliani, A., Gheorghe, M., Castro-Mondragon, J.A., Van Der Lee, R., Bessy, A., Chèneby, J., Kulkarni, S.R., Tan, G., et al. (2018). JASPAR 2018: Update of the open-access database of transcription factor binding profiles and its web framework. Nucleic Acids Res.

Kirkham, B.W., Lassere, M.N., Edmonds, J.P., Juhasz, K.M., Bird, P.A., Lee, C.S., Shnier, R., and Portek, I.J. (2006). Synovial membrane cytokine expression is predictive of joint damage progression in rheumatoid arthritis: a two-year prospective study (the DAMAGE study cohort). Arthritis Rheum. 54, 1122–1131.

Koga, T., Yamasaki, S., Migita, K., Kita, J., Okada, A., Kawashiri, S., Iwamoto, N., Tamai, M., Arima, K., Origuchi, T., et al. (2011). Post-transcriptional regulation of IL-6 production by Zc3h12a in fibroblast-like synovial cells. Clin. Exp. Rheumatol. 29, 906–912.

Koshy, P.J., Henderson, N., Logan, C., Life, P.F., Cawston, T.E., and Rowan, A.D. (2002). Interleukin 17 induces cartilage collagen breakdown: novel synergistic effects in combination with proinflammatory cytokines. Ann. Rheum. Dis. 61, 704–713.

Kotake, S., Udagawa, N., Takahashi, N., Matsuzaki, K., Itoh, K., Ishiyama, S., Saito, S., Inoue, K., Kamatani, N., Gillespie, M.T., et al. (1999). IL-17 in synovial fluids from patients with rheumatoid arthritis is a potent stimulator of osteoclastogenesis. J. Clin. Invest. 103, 1345–1352.

Kühnemuth, B., Mühlberg, L., Schipper, M., Griesmann, H., Neesse, A., Milosevic, N., Wissniowski, T., Buchholz, M., Gress, T.M., and Michl, P. (2013). CUX1 modulates polarization of tumor-associated macrophages by antagonizing NF-κB signaling. Oncogene 34, 177–187.

Liberzon, A., Birger, C., Thorvaldsdóttir, H., Ghandi, M., Mesirov, J.P., and Tamayo, P. (2015). The Molecular Signatures Database Hallmark Gene Set Collection. Cell Syst. 1, 417–425.

Lubberts, E. (2015). The IL-23-IL-17 axis in inflammatory arthritis. Nat. Rev. Rheumatol. 11, 415–429.

Miossec, P. (2017). Update on interleukin-17: a role in the pathogenesis of inflammatory arthritis and implication for clinical practice. RMD Open 3, e000284.

Miossec, P., and Kolls, J.K. (2012). Targeting IL-17 and TH17 cells in chronic inflammation. Nat. Rev. Drug Discov. 11, 763–776.

Mix, K.S., McMahon, K., McMorrow, J.P., Walkenhorst, D.E., Smyth, A.M., Petrella, B.L., Gogarty, M., Fearon, U., Veale, D., Attur, M.G., et al. (2012). Orphan nuclear receptor NR4A2 induces synoviocyte proliferation, invasion, and matrix metalloproteinase 13 transcription. Arthritis Rheum. 64, 2126–2136.

Monin, L., and Gaffen, S.L. (2018). Interleukin 17 Family Cytokines: Signaling Mechanisms, Biological Activities, and Therapeutic Implications. Cold Spring Harb. Perspect. Biol. 10.

Moran, E.M., Mullan, R., McCormick, J., Connolly, M., Sullivan, O., Fitzgerald, O., Bresnihan, B., Veale, D.J., and Fearon, U. (2009). Human rheumatoid arthritis tissue production of IL-17A drives matrix and cartilage degradation: synergy with tumour necrosis factor-alpha, Oncostatin M and response to biologic therapies. Arthritis Res. Ther. 11, R113.

Mulherin, D., Fitzgerald, O., and Bresnihan, B. (1996). Synovial tissue macrophage populations and articular damage in rheumatoid arthritis. Arthritis Rheum. 39, 115–124.

Muromoto, R., Hirao, T., Tawa, K., Hirashima, K., Kon, S., Kitai, Y., and Matsuda, T. (2016). IL-17A plays a central role in the expression of psoriasis signature genes through the induction of IκB-ζ in keratinocytes. Int. Immunol. 28, 443–452.

Nguyen, H.N., Noss, E.H., Mizoguchi, F., Huppertz, C., Wei, K.S., Watts, G.F.M., and Brenner, M.B. (2017). Autocrine Loop Involving IL-6 Family Member LIF, LIF Receptor, and STAT4 Drives Sustained Fibroblast Production of Inflammatory Mediators. Immunity 46, 220–232.

Onishi, R.M., and Gaffen, S.L. (2010). Interleukin-17 and its target genes: mechanisms of interleukin-17 function in disease. Immunology 129, 311–321.

Panopoulos, A.D., and Watowich, S.S. (2008). Granulocyte colony-stimulating factor: molecular mechanisms of action during steady state and “emergency” hematopoiesis. Cytokine 42, 277–288.

Parrish, W.R., Byers, B.A., Su, D., Geesin, J., Herzberg, U., Wadsworth, S., Bendele, A., and Story, B. (2017). Intra-articular therapy with recombinant human GDF5 arrests disease progression and stimulates cartilage repair in the rat medial meniscus transection (MMT) model of osteoarthritis. Osteoarthr. Cartil. 25, 554–560.

Parsonage, G., Filer, A., Bik, M., Hardie, D., Lax, S., Howlett, K., Church, L.D., Raza, K., Wong, S.-H., Trebilcock, E., et al. (2008). Prolonged, granulocyte--macrophage colony-stimulating factor-dependent, neutrophil survival following rheumatoid synovial fibroblast activation by IL-17 and TNFalpha. Arthritis Res. Ther. 10, R47.

R.G., L., B.E., E., M., L., K., R., C.E.M., G., K., P., L., P., H., N., L., S., B., S., et al. (2014). Secukinumab in plaque psoriasis - Results of two phase 3 trials. N. Engl. J. Med.

Raychaudhuri, S., Stuart, J.M., and Altman, R.B. (2000). Principal components analysis to summarize microarray experiments: application to sporulation time series. Pac. Symp. Biocomput. 38, 455–466.

Ritchie, M.E., Phipson, B., Wu, D., Hu, Y., Law, C.W., Shi, W., and Smyth, G.K. (2015). limma powers differential expression analyses for {RNA-sequencing} and microarray studies. Nucleic Acids Res. 43, e47.

Roelofs, A.J., Zupan, J., Riemen, A.H.K., Kania, K., Ansboro, S., White, N., Clark, S.M., and De Bari, C. (2017). Joint morphogenetic cells in the adult mammalian synovium. Nat. Commun. 8, 15040.

Ruddy, M.J., Wong, G.C., Liu, X.K., Yamamoto, H., and others (2004). Functional cooperation between interleukin-17 and tumor necrosis factor-α is mediated by CCAAT/enhancer-binding protein family members. J. Biol.

Shen, F., Ruddy, M.J., Plamondon, P., and Gaffen, S.L. (2005). Cytokines link osteoblasts and inflammation: microarray analysis of interleukin-17- and TNF-alpha-induced genes in bone cells. J. Leukoc. Biol. 77, 388–399.

Smolen, J.S., and Aletaha, D. (2015). Rheumatoid arthritis therapy reappraisal: strategies, opportunities and challenges. Nat. Rev. Rheumatol. 11, 276–289.

Yamamoto, M., Yamazaki, S., Uematsu, S., Sato, S., Hemmi, H., Hoshino, K., Kaisho, T., Kuwata, H., Takeuchi, O., Takeshige, K., et al. (2004). Regulation of Toll/IL-1-receptor-mediated gene expression by the inducible nuclear protein IkappaBzeta. Nature 430, 218–222.

Yamazaki, S., Matsuo, S., Muta, T., Yamamoto, M., Akira, S., and Takeshige, K. (2008). Gene-specific requirement of a nuclear protein, IkappaB-zeta, for promoter association of inflammatory transcription regulators. J. Biol. Chem. 283, 32404–32411.

Zhang, F., Wei, K., Slowikowski, K., Fonseka, C.Y., Rao, D.A., Kelly, S., Goodman, S.M., Tabechian, D., Hughes, L.B., Salomon-Escoto, K., et al. (2018). Defining Inflammatory Cell States in Rheumatoid Arthritis Joint Synovial Tissues by Integrating Single-cell Transcriptomics and Mass Cytometry. BioRxiv.

Zhang, Q., Lenardo, M.J., and Baltimore, D. (2017). 30 Years of NF-κB: A Blossoming of Relevance to Human Pathobiology. Cell 168, 37–57.

Zrioual, S., Ecochard, R., Tournadre, A., Lenief, V., Cazalis, M.-A., and Miossec, P. (2009). Genome-wide comparison between IL-17A- and IL-17F-induced effects in human rheumatoid arthritis synoviocytes. J. Immunol. 182, 3112–3120.

Zvaifler, N.J. (1973). The immunopathology of joint inflammation in rheumatoid arthritis. Adv. Immunol. 16, 265–336.

